# Single-cell transcriptional dynamics in a living vertebrate

**DOI:** 10.1101/2024.01.03.574108

**Authors:** Elizabeth Eck, Bruno Moretti, Brandon H. Schlomann, Jordão Bragantini, Merlin Lange, Xiang Zhao, Shruthi VijayKumar, Guillaume Valentin, Cristina Loureiro, Pablo Perez Franco, Chloé Jollivet, Baldemar Motomochi, Loïc A. Royer, Andrew C. Oates, Hernan G. Garcia

## Abstract

The ability to quantify transcriptional dynamics in individual cells via live imaging has revolutionized our understanding of gene regulation. However, such measurements are lacking in the context of vertebrate embryos. We addressed this deficit by applying MS2-MCP mRNA labeling to the quantification of transcription in zebrafish, a model vertebrate. We developed a platform of transgenic organisms, light sheet fluorescence microscopy, and optimized image analysis that enables visualization and quantification of MS2 reporters. With these tools, we obtained single-cell, real-time measurements of transcriptional dynamics of the segmentation clock. Our measurements challenge the traditional view of smooth clock oscillations and instead suggest a model of discrete transcriptional bursts that are organized in space and time. Together, these results highlight how measuring single-cell transcriptional activity in the context of vertebrate organisms can reveal unexpected features of gene regulation and how this data can fuel the dialogue between theory and experiment.

## Main text

Development is driven by highly dynamic and coordinated gene regulatory programs. Rapid switches (Afzal and Krumlauf 2022), transient excursions (Bothma et al. 2014), and oscillations (Kageyama et al. 2008; Palmeirim et al. 1997) in gene expression are commonplace across the tree of life. Innovations over the last few years have resulted in an explosion of our ability to map the gene regulatory networks dictating developmental dynamics. However, predictively understanding the links between the transcriptional dynamics, cell fate decisions, and tissue patterning encoded by these networks remains an open, fundamental problem in developmental biology (Garcia et al. 2020).

The MS2-MCP system is a widely used method for precise, quantitative measurements of real-time transcriptional dynamics of individual genes (Bertrand et al. 1998; Golding et al. 2005; Chao et al. 2008; Larson et al. 2011; Garcia et al. 2013; Lucas et al. 2013) (Fig. 1A). In this system, a transgene is tagged with multiple copies of a stem loop sequence from bacteriophage MS2 (“MS2”), whose cognate binding partner (“MCP”, MS2 coat protein) is fused to a fluorescent protein. Upon mRNA synthesis, MCP binds the MS2 loops within the nascent transcript, producing a region of enhanced fluorophore concentration (a “spot”) that can be detected with a fluorescence microscope (Fig. 1A). The intensity of the resulting fluorescent spot is proportional to the number of RNA polymerase molecules actively transcribing the gene.

**Figure 1:**
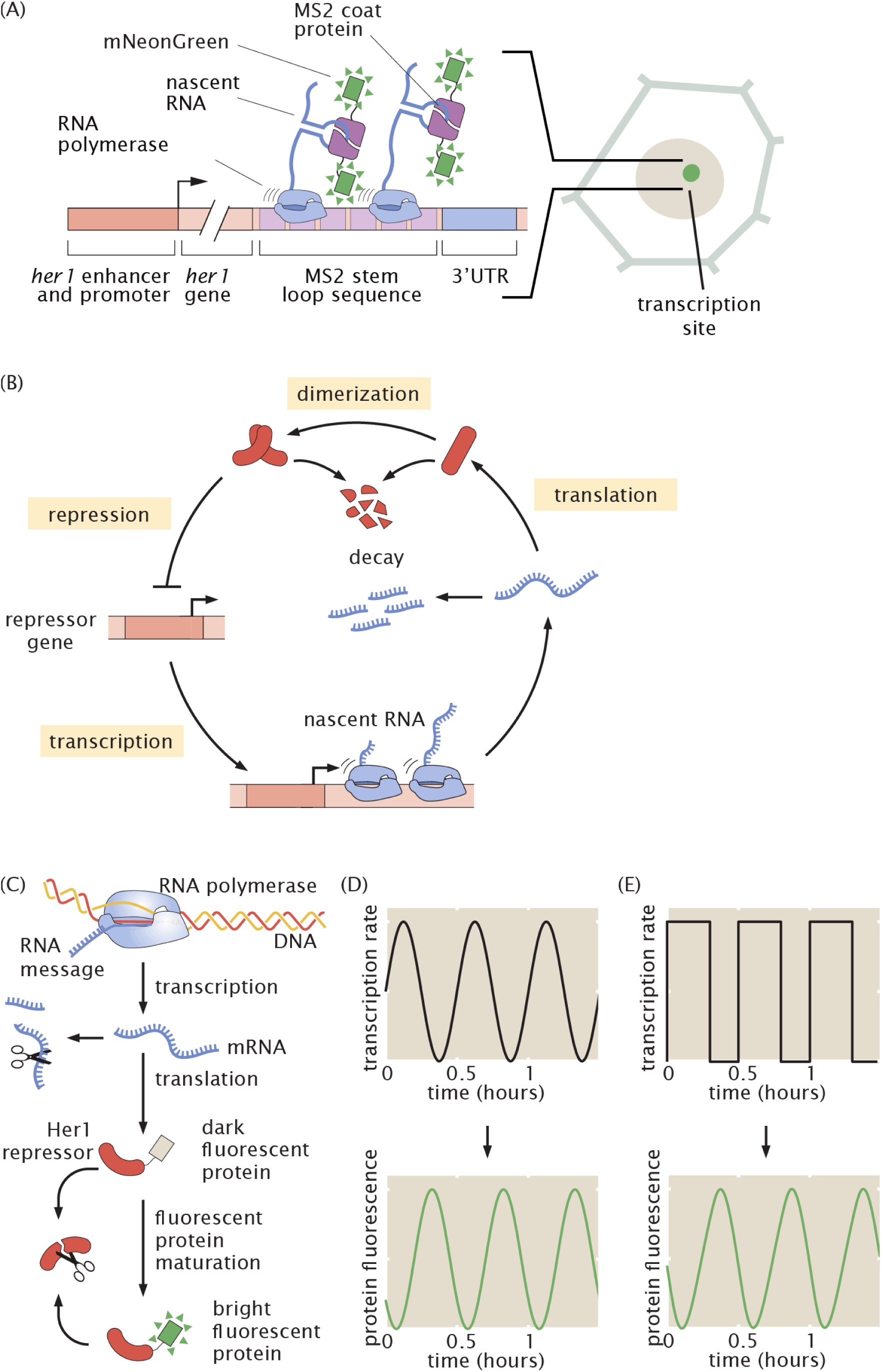
Protein reporters cannot distinguish between continuous and discrete modes of transcriptional regulation of the segmentation clock. (A) Schematic of the MS2 system, in which pre-formed fluorescent proteins (mNeonGreen) fused to MS2 coat protein bind to nascent mRNA strands via a stem loop-coat protein interaction. Fluorescence accumulates at sites of nascent transcript formation. In this way, the MS2 system gives a near-real time readout of transcriptional activity. In this work, we tag a transgene containing the segmentation clock gene *her1* and its regulatory region. (B) Oscillations are generated via auto-repression of segmentation clock genes such as *her1*. (C) Schematic of the central dogma showing the steps that lead to low-pass filtering between transcriptional and protein dynamics. (D, E) The low-pass filtering effect of product accumulation and decay leads to smooth protein oscillations that can be generated by dissimilar transcriptional dynamics ranging from smooth modulation (D) to discrete pulses (E).

To date, precise, quantitative measurements of transcriptional dynamics via the MS2-MCP system have been limited to single cells in culture, tissue samples, invertebrate model organisms, such as flies and worms, and plants (Garcia et al. 2013; Lammers, Kim, et al. 2020; Wildner et al. 2023; Lee, Shin, and Kimble 2019; Corrigan et al. 2016; Alamos et al. 2021; Park et al. 2014). In these contexts, the MS2-MCP system has transformed our understanding of gene expression by revealing highly dynamic gene regulatory programs and novel molecular mechanisms of transcriptional regulation (Lim 2018). However, similar measurements have yet to be achieved in a living, intact vertebrate. Given its established genetic toolkit and amenability to live imaging, the zebrafish is a prime candidate for exploring single-cell transcriptional dynamics in vertebrates. While MS2 has been used in zebrafish to study mRNA localization, previous implementations have faced issues of MCP aggregation that prevented quantification of transcription (Campbell et al. 2015; Fedder-Semmes and Appel 2021; Costa et al. 2020; Zhang et al. 2020).

Further, even if the genetic implementation of MS2 in vertebrates such as zebrafish was optimized, large challenges to using this technology to quantify transcriptional dynamics in intact vertebrate animals remain. For example, compared to flies or worms, zebrafish embryos are larger, denser, and exhibit greater motion during development, posing challenges in both image acquisition and analysis. Due to fast cell motion during tailbud elongation (50 μm/h is a typical speed; see (Lawton et al. 2013)), fields of view hundreds of micrometers in length are required to track groups of cells for multiple hours. Simultaneously, one must be able to detect signals coming from diffraction-limited spots made of only tens to hundreds of fluorophores. Large fields of view, over time, result in large (here, ∼1 TB) datasets. The large size of the data, together with dense nuclei and strong, heterogeneous fluorescence backgrounds, complicate cell tracking and MS2 spot quantification. Thus, the full power of the MS2 system to quantify transcriptional dynamics has yet to be realized in zebrafish or any other vertebrate animal for molecular biology, imaging, and data processing reasons. As many aspects of vertebrate development and physiology fundamentally differ from invertebrates, entire fields of questions concerning transcriptional dynamics—including questions of direct biomedical relevance—remain unanswerable.

Here, we report the establishment of the MS2 system in transgenic zebrafish together with the development of an allied platform of light sheet fluorescence microscopy and computational image analysis to quantify transcriptional dynamics of MS2 reporters in complex and dynamical vertebrate embryos. To make this possible, we generated a new transgenic zebrafish line expressing a reengineered MCP that avoids protein aggregation. Further, to enable fast, large volume imaging, we used light sheet fluorescence microscopy and demonstrated that this method can retain the sensitivity required to detect subcellular MS2 spots while simultaneously imaging tens of thousands of cells in intact tissue. To enable single cell tracking and spot quantification in large datasets with strong background, we developed a computational image analysis pipeline that combines parallelized chunk-based file handling, GPU-accelerated image processing, and deep learning, achieving fast and accurate measurements of transcriptional dynamics in single cells.

We applied this platform to a paradigmatic example of vertebrate-specific gene expression dynamics: the segmentation clock. In vertebrate embryo development, somites—morphological segments that prefigure the bones and muscles of the adult—are formed rhythmically and sequentially at the posterior end of the elongating body axis from the presomitic mesoderm (Oates, Morelli, and Ares 2012; Palmeirim et al. 1997). This rhythmic specification is dictated by a highly conserved biological clock consisting of an oscillatory gene regulatory network (Oates, Morelli, and Ares 2012) which is rooted in an auto-repression motif (Fig. 1B). Protein reporters have revealed insights into the dynamics of these oscillators, revealing smooth, sinusoidal oscillations at both the tissue scale (Soroldoni et al. 2014; Webb et al. 2016; Zinani et al. 2020) and in single cells (Delaune et al. 2012; Shih et al. 2015; Simsek et al. 2023; Venzin et al. 2023; Webb et al. 2016; Rohde et al. 2024). However, how oscillations are generated at the transcriptional level remains unknown. Due to the low-pass filtering effect of RNA and protein accumulation and degradation (Fig. 1C), high frequency dynamics are smoothed out at the protein level, making these transcriptional dynamics invisible to measurements afforded by the protein reporters available to date. For example, single-cell transcription rates could mirror protein levels, oscillating in a smooth, sinusoidal fashion (Fig. 1D), or they could be sharp and discrete (Fig. 1E). Both scenarios would produce smooth protein oscillations.

At the tissue scale, our transcriptional measurements recapitulate the known global picture of somitogenesis: the expression of the core clock gene *her1* oscillates over time and travels in a wave-like pattern from the tailbud to the forming somites. However, at the single-cell level we discovered that, in contrast to the smooth cycles of protein levels observed with fluorescent protein fusions in all vertebrate systems studied to date (Diaz-Cuadros et al. 2020; Yoshioka-Kobayashi et al. 2020; Tsiairis and Aulehla 2016; Delaune et al. 2012; Rohde et al. 2024), transcription rate is modulated in a sharp, discrete manner, resulting in bursts of transcription. While these bursts are largely periodic, they also exhibit a degree of stochasticity. Simulations of the protein oscillations predicted to occur based on these transcriptional dynamics are regular and sinusoidal, indicating a mechanism of robust oscillations that is compatible with this stochastic, pulse-like transcription. To begin to explore how regular oscillations arise from transcriptional bursting, we developed a minimal mathematical model that extends the canonical two-state bursting model (Lammers, Kim, et al. 2020) to include auto-repressive feedback. Analysis of waiting time distributions for the active and inactive transcriptional intervals provides evidence for a molecular mechanism of *her1* action consisting in either the regulation of burst amplitude or the regulation of a combination of frequency and duration.

Altogether, this work provides the tools necessary for studying transcriptional dynamics in zebrafish to the greater community and presents new measurements that challenge the existing paradigm of somitogenesis. Further, the pipeline presented here constitutes a framework that can be used to launch quantitative live imaging and theoretical dissection of transcriptional dynamics in other vertebrate systems.

## Results

### Implementing MS2 in zebrafish embryos to quantify transcriptional dynamics of the segmentation clock

While expression of transgenes via transient injection of one-cell stage embryos is common in zebrafish (Campbell et al. 2015), this approach is not ideal for reproducible, quantitative measurements of transcription due to sources of variation such as mosaicism and positional effects (Stuart et al. 1990; Roberts et al. 2014). Therefore, we generated two stable transgenic zebrafish lines via I-SceI meganuclease transgenesis (Soroldoni, Hogan, and Oates 2009): one carrying a reporter construct containing MS2 stem loop sequences inserted downstream of the *her1* regulatory region, dubbed “*her1*-MS2”, and another carrying an MCP-mNeonGreen (herein “MCP”) fusion driven by a ubiquitin promoter (Mosimann et al. 2011) (Fig. 2A-B, see the “Zebrafish” section of the Methods). The *her1*-MS2 construct was built off of previously established *her1*-Venus fusion protein transgene construct, with the fluorescent protein switched for the MS2 stem loops (Soroldoni et al. 2014). Our approach differs from previous works on the MS2 system in zebrafish, which used transient injection of reporter constructs containing the MS2 loops (Campbell et al. 2015; Fedder-Semmes and Appel 2021; Costa et al. 2020; Zhang et al. 2020).

**Figure 2:**
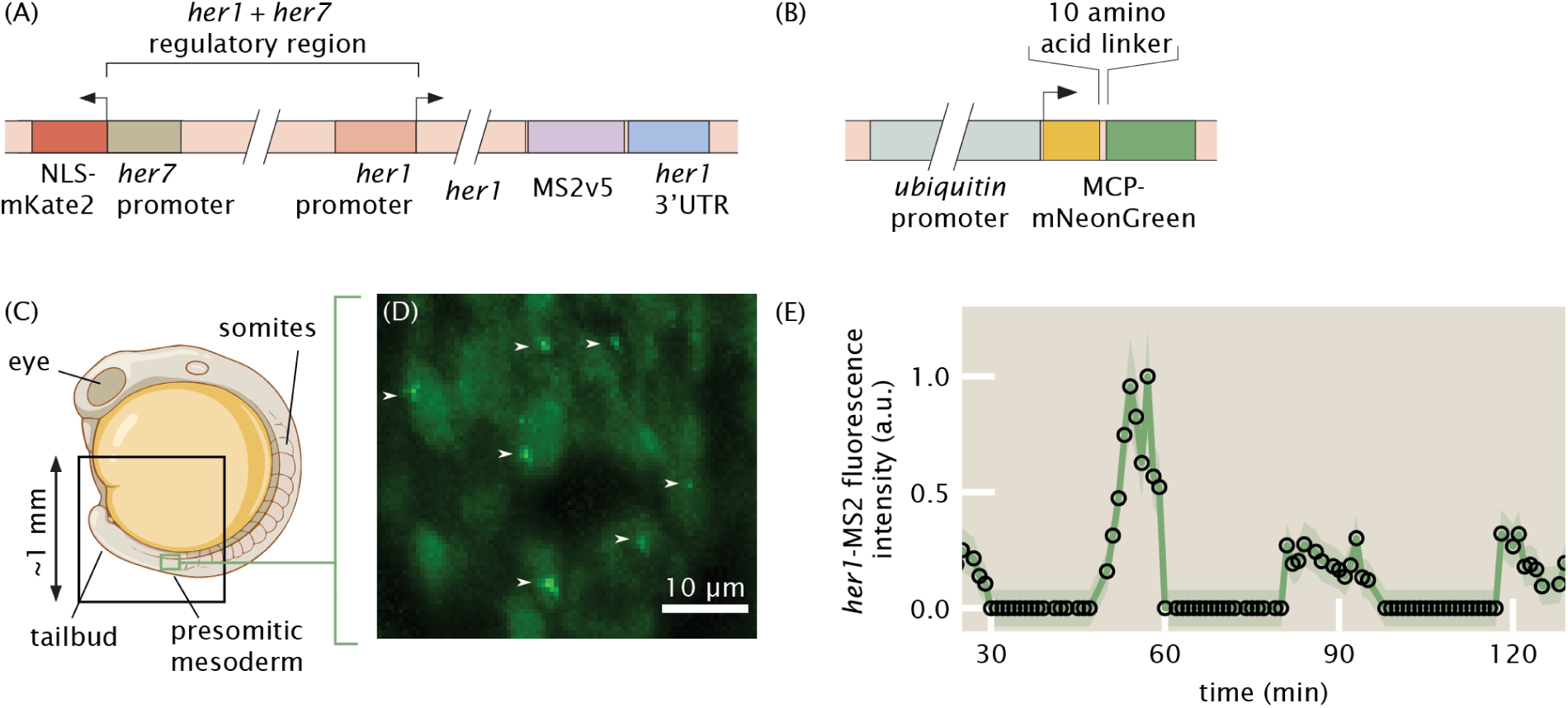
The MS2 system can be used to visualize transcriptional dynamics of the zebrafish segmentation clock. (A)-(B) Details of the transgenic MS2 (A) and MCP (B) constructs. Stable transgenic lines were made using I-SceI meganuclease transgenesis. The MS2 construct contains an mKate2 marker for ease of screening transgenic zebrafish. (C) Geometry of a zebrafish embryo. Black box represents the approximate field of view of the imaging experiment. Green box represents the approximate field of view of the zoomed-in area shown in (D). (D) Maximum intensity projection over 50 micrometers of the signal from MCP-mNeonGreen, *her1*-MS2 zebrafish showing several fluorescent spots (white arrows) against the hazy background MCP expression in individual nuclei. (E) Example *her1*-MS2 trace obtained for an individual cell within the presomitic mesoderm showing oscillations. Error bars are reported by the shaded area and reflect uncertainty due to background fluctuations and spot detection (see the “Spot analysis” section of the Methods). Panel (C) is adapted from BioRender.

We also made modifications to the MCP construct. Specifically, previous work used tandem dimer MCP with a nuclear localization signal (Campbell et al. 2015) and saw unwanted aggregation of the coat protein. We suspected that this aggregation was due to tandem dimer MCP and therefore used regular MCP. Further, while the nuclear localization signal has proven useful for keeping background levels low when monitoring transcript localization in the cytoplasm (Forrest and Gavis 2003), it can lead to overly high background levels in the nucleus that interfere with transcription measurements. Therefore, we did not include a nuclear localization signal in our construct.

We used I-SceI transgenesis because it is highly efficient and typically generates single insertion events (Soroldoni, Hogan, and Oates 2009). Single insertion events are ideal here because multiple insertion events could result in multiple MS2 spots, which would complicate our analyses. However, I-SceI transgenesis can insert multiple copies of the transgene at the same site (Soroldoni, Hogan, and Oates 2009). Indeed, based on qPCR we measured that our line carries 4 ± 2 copies of the *her1*-MS2 reporter (Methods). These multiple copies raise two potential issues. First, multiple copies of the *her1* promoter might activate asynchronously, which would produce complicated MS2 signals. As we demonstrate below, we observe largely periodic signals, indicating that the multiple copies are acting in unison to a high degree. We elaborate on this point further in the Discussion section.

A second potential issue is that, since our construct contains the full *her1* coding sequence, it is likely that extra functional copies of Her1 are produced, altering clock function and producing developmental defects. To address this potential issue, we performed extensive characterization of segmentation clock patterning and somite formation in our transgenic lines. For both MS2 and MCP zebrafish lines, heterozygous animals showed normal patterns of *her1* expression (Supplemental Fig. 1A-D) and minimal somite boundary formation defects consistent with previous Her1 protein reporter lines (Soroldoni et al. 2014; Delaune et al. 2012) (Supplemental Fig. 1F), indicating healthy somitogenesis. Further, in situ hybridization against the MS2 sequence mirrored the patterns of *her1* expression (Supplemental Fig. 1E), demonstrating that our construct faithfully reports on *her1* activity.

To assess the feasibility of detecting *her1*-MS2 spots, we imaged embryos on multiple types of fluorescence microscopes. To visualize nuclei, we crossed MCP-mNeonGreen fish with a line carrying h2b-mScarlet (O’Brown, Megason, and Gu 2019) (see the “Zebrafish” section of the Methods). In *her1*-MS2/+; MCP-mNeonGreen/+; h2b-mScarlet/+ animals, green fluorescent puncta representing nascent *her1*-MS2 transcripts were readily detectable using both light sheet (Fig. 2C-D, Supplemental Fig. 2A-B) and laser-scanning confocal (Supplemental Fig. 2C-D) fluorescence microscopes. In control experiments with embryos expressing only MCP-mNeonGreen and no *her1*-MS2, fluorescent puncta were detected (Supplemental Fig. 2B).

In initial time lapse imaging experiments on a laser scanning confocal (Zeiss 880 with AiryScan), we found that achieving even a barely-detectable MS2 signal with 1 minute time resolution (required to capture substructure within the oscillations, whose period is ∼30 min) resulted in a small field of view of 100×50×20 μm^3^, or approximately 50 nuclei. To image the full tissue would require a field of view of approximately 500×500×400 μm^3^. Due to significant cell motion during somitogenesis (Lawton et al. 2013), we found that manual adjustment of the microscope stage was required every few minutes to keep the same cells in the field of view (<u>Movie 1</u>). To capture multiple oscillations, this constant adjustment must be done for multiple hours and the resulting movies registered together, making this approach impractical. Consequently, we exclusively used light sheet fluorescence microscopy, which enabled us to capture the full tissue containing somites, presomitic mesoderm, and tailbud, with 1 minute time resolution and acceptable signal-to-noise ratios (Royer et al. 2016) (see the “Light sheet imaging” section of the Methods).

We built a computational image analysis pipeline (Supplemental Fig. 3A, B) to extract quantitative measurements of transcriptional dynamics from our images. As elaborated in the “Spot analysis” section of the Methods, the large size (∼1 TB) of the image datasets and the strong and heterogeneous background signal posed new challenges to the analysis of MS2 data. Specifically, the large size of the data called for out-of-memory computing schemes and speed optimization of every step of the pipeline, while the strong background required sophisticated algorithms for spot classification and localization. Overall, these challenges, which were new to the analysis of MS2 data in general, necessitated the development of specialized software. Our pipeline tracks nuclei and identifies spots in parallel, then spots are assigned to nuclei and the sequences of MS2 spot intensities over time—”traces”—are constructed. Initial examination of traces revealed clear oscillations (Fig. 2E).

We performed extensive characterization of the pipeline’s accuracy (see the “Spot analysis” section of the Methods, Supplemental Fig. 3C-H). Through comparison with 26 manually obtained traces, we found that our automated 3D nuclear tracking was approximately 80% accurate (21/26 nuclear trajectories showed no errors), with an error rate of around 1 nuclear tracking error per 5 nuclear trajectories, or on average 1 error per 500 minutes (see also <u>Movie 2</u>). Examples of tracking errors include swapping nuclear identities between neighboring cells, grouping two nuclei into one, and splitting one nucleus into two (Supplemental Fig. 3G).

Through comparison with manually curated traces, where spot detection was done entirely by human visual inspection, we found that spot detection was extremely accurate. Specifically, we found a false positive rate of 4% and a false negative rate of 8% (Supplemental Fig. 3H, I). When both manual traces and pipeline traces detected a spot, the resulting spot intensities were highly correlated with an *R*^2^ of 0.91 (Supplemental Fig. 3I). Through analysis of synthetic images containing simulated spots, we found that our spot fluorescence quantification algorithm was also highly accurate, with typical errors of around 5% (Supplemental Fig. 4, Methods). As our MCP background was expressed quite heterogeneously throughout the tissue, it is possible that in some cells MCP levels were low enough to not saturate the MS2 stem loops, introducing a background-dependent signal (Wu, Chao, and Singer 2012). However, we observed only a very weak correlation (*R*^2^=0.26) between the fluorescence background of spots and their (background subtracted) signal (Supplemental Fig. 3E), indicative of MCP saturation and reliable measurements of absolute fluorescence intensity.

Taken together, these results demonstrate that we successfully implemented the MS2 system in zebrafish for quantifying *her1* transcriptional dynamics. We then turned to studying the behavior of *her1* transcription at tissue and single-cell scales.

### *her1* transcriptional dynamics recapitulate protein oscillations and wave patterns at the tissue scale

We first assessed how the tissue-scale dynamics of the segmentation clock at the transcriptional level compare to the measurements obtained from Her1/Her7 protein dynamics reported by fluorescent protein fusions (Soroldoni et al. 2014; Delaune et al. 2012; Shih et al. 2015; Webb et al. 2016; Zinani et al. 2020; Simsek et al. 2023; Venzin et al. 2023; Rohde et al. 2024). Previous protein measurements revealed collective oscillations whose period increases as cells approach somite formation, resulting in a phase wave that travels from posterior to anterior, arresting at the formation of a new somite (Soroldoni et al. 2014; Delaune et al. 2012). The oscillation period varies with temperature, being approximately 30 minutes at 28 C.

While the dynamics of protein reporters can be directly visualized in movies (Soroldoni et al. 2014; Delaune et al. 2012), MS2 spots are too small and dim to be picked up in raw 3D renderings or projections of the full tissue. Therefore, to visualize transcriptional dynamics, we false colored nuclei in proportion to their spot intensity (Fig. 3A-C, <u>Movie 4</u>, <u>Movie 5</u>). The resulting movies showed oscillations and waves similar to movies of fluorescent proteins, qualitatively recapitulating previous protein-level dynamic measurements, albeit with a higher prevalence of noise and high frequency flashing.

**Figure 3:**
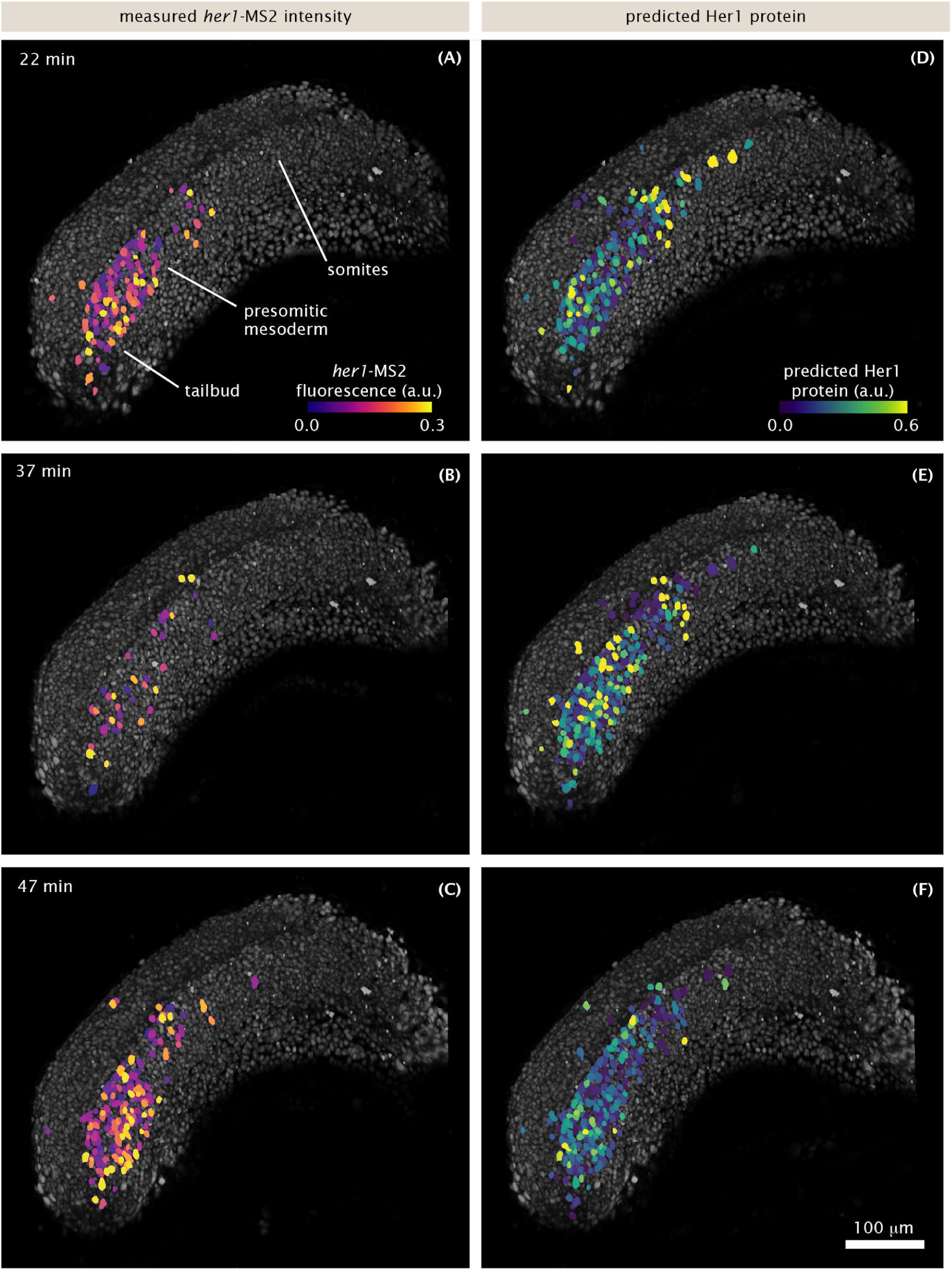
Transcriptional dynamics at the tissue scale recapitulate known features of protein patterns. Snapshots of 3D renderings of the presomitic mesoderm and tailbud during somitogenesis. (A-C): nuclei (gray) are false-colored in proportion to the intensity of their MS2 spots. (D-F): same as (A-C) but nuclei are colored in proportion to the predicted Her1 protein level, given the MS2 signal in (A-C) (Methods). See also <u>Movie 4</u>, <u>Movie 5</u>, and <u>Movie 6</u>. Scale bar: 100 μm. Time is measured from the start of image acquisition, which corresponds to approximately the 15 somite stage.

To validate our construct more directly against previous protein measurements, we compared the protein dynamics predicted by our transcriptional data to actual protein-level measurements of Her1 dynamics. Using a linear production/decay model (based on Fig. 1C, Methods) and our MS2 traces as input, we generated visualizations of the predicted Her1 protein concentrations (Fig. 3D-F, <u>Movie 6</u>). The predicted protein patterns are smoother than the MS2 patterns, showing gradual waves traveling from posterior to anterior that resemble previous measurements (Soroldoni et al. 2014; Delaune et al. 2012). Thus, once protein production and decay are taken into account, our transcriptional reporter faithfully recapitulates the behavior of established protein reporters at a qualitative level.

To quantitatively validate our reporter, we captured transcription and protein spatiotemporal patterns in kymographs (Fig. 4). To make this possible, we defined an anterior-posterior axis with a combination of manual labeling and spline interpolation (Fig. 4A, <u>Movie 7</u>, Methods). We grouped cells into equally-spaced bins along this axis, summed the spot intensities in each bin and plotted them over time. Based on previous measurements (Soroldoni et al. 2014), we expected the kymographs to resemble the schematic in Fig. 4B (Methods), which we briefly describe here. In these kymographs, with time increasing downwards and anterior to the right, horizontal stripes correspond to the stationary oscillations that occur in the tailbud, while diagonal stripes correspond to waves that travel towards the anterior and arrest as somites form. If one picks a point in absolute space and follows the oscillations over time, the oscillation period increases. We hypothesized that these features, which were identified in kymographs of Her1 protein (Soroldoni et al. 2014), should be apparent in the kymograph of *her1* transcriptional dynamics.

**Figure 4:**
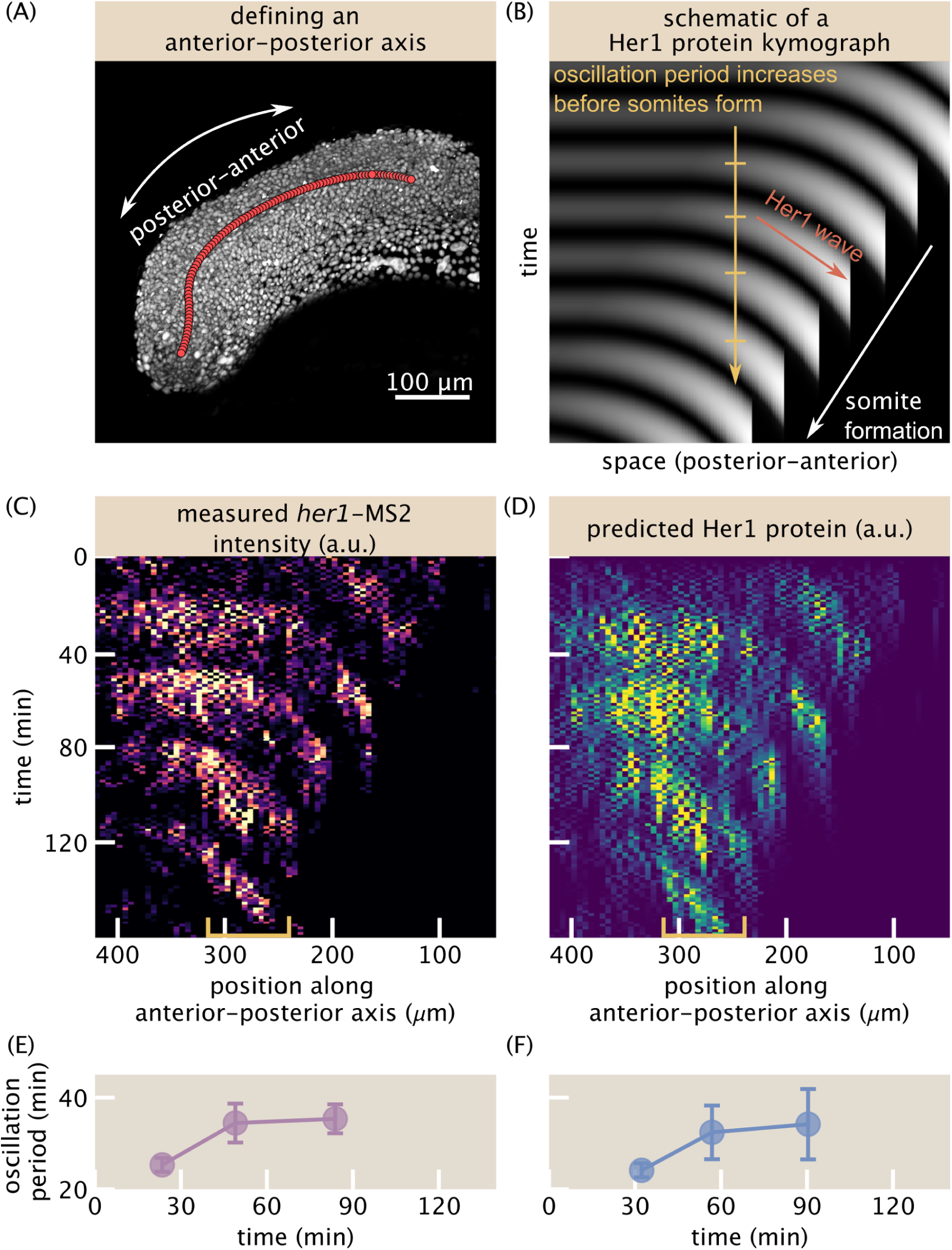
Tissue-scale analysis reveals transcriptional oscillations and wave-like patterns. (A) 3D rendering of nuclei (gray) illustrating the anterior-posterior axis (red circles). The axis is defined coarsely every 50 time points, roughly following the notochord, and then spline interpolated in both space and time (see the “Anterior-posterior axis” section of the Methods). See also <u>Movie 7</u>. (B) Schematic of a Her1 protein kymograph highlighting the known features. Oscillation period increases as cells approach somite formation, which produces a traveling wave pattern. (C-D) Kymographs of total *her1*-MS2 (C) spot intensity and predicted Her1 protein (D). Distance is defined along the anterior-posterior axis as in (A). Spot anterior-posterior positions are binned, with bin size = 6.5 μm, and the sum of all intensities in each bin is plotted for each time point. (E)-(F) Measurement of the increase in oscillation period as cells approach somite formation for the raw *her1*-MS2 kymograph (E) and the predicted Her1 protein kymograph (F). The regions used to compute the period are noted by the yellow brackets in panels (C) and (D).

The resulting MS2 kymograph (Fig. 4C) shows clear, regular oscillations throughout the tissue, with periods of approximately 30 min, consistent with protein measurements (Soroldoni et al. 2014). Further, the left-down sloping of the stripes indicates the traveling phase wave from posterior to anterior, with a speed of approximately 100 μm/h, also consistent with previous measurements (Soroldoni et al. 2014). The predicted protein kymograph (Fig. 4D) largely mirrors the MS2 kymograph, though the protein kymograph is visibly smoother and phase shifted with respect to the MS2 one, as expected from the time lag between transcription and translation. Quantitatively, we observed an increase in the oscillation period in the kymograph of approximately 40% prior to somite formation for both MS2 and predicted protein signals (Fig. 4E-F).

All together, these observations demonstrate that, at the tissue scale, the transcriptional dynamics of the segmentation clock largely mirror protein dynamics, albeit with transcriptional activity appearing noisier than accumulated protein levels.

### Single-cell transcriptional oscillations comprise sharp, quasi-periodic bursts

Exploiting the unique power of the MS2 system, we next investigated transcriptional dynamics in single cells in a set of 101 traces that were manually checked and corrected as needed. Individual MS2 traces (Fig. 5A, green lines; Supplemental Fig. 5) show clear periodicity. However, in contrast to previously measured smooth protein oscillations (Delaune et al. 2012; Soroldoni et al. 2014; Rohde et al. 2024), our observed transcriptional dynamics are strikingly sharp and discrete. Further, we observed examples of multiple, distinct transcriptional pulses in rapid succession, which we refer to as bursts (Fig. 5A, black arrows). We visually inspected these bursts and confirmed the rapid disappearance and reappearance of the MS2 spot (Fig. 5B, C). Remarkably, even for these cases with a higher frequency of transcriptional bursting, predicted protein traces for single cells are smooth, roughly sinusoidal, and regular in period, similar to measured protein traces (Fig. 5A, cyan lines; Methods) (Delaune et al. 2012; Rohde et al. 2024). With a conservative burst-calling algorithm (Methods), we measured that 8 ± 3% (mean ± std. dev. from bootstrapping over cells) of protein oscillations are generated by 2 or more transcriptional bursts (Fig. 5E, Methods). Comparing the periods of transcriptional bursts and the predicted protein oscillations, we found that the predicted protein oscillations were more regular than the transcriptional bursts, having a narrower, more peaked distribution (Fig. 5F). Thus, our measurements of transcriptional bursts in the segmentation clock are consistent with the observation of regular protein oscillations, even without invoking additional levels of post-transcriptional regulation.

**Figure 5:**
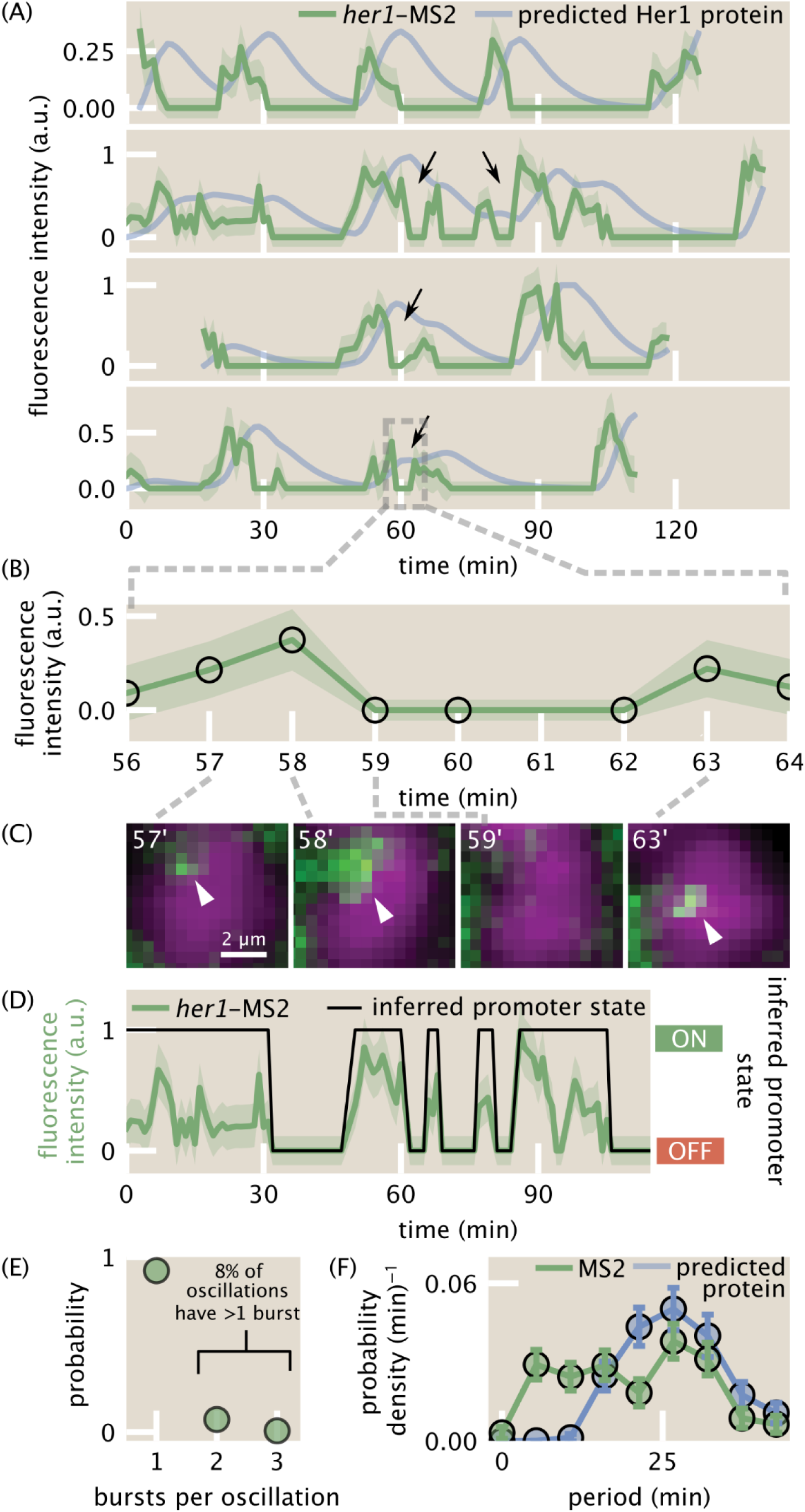
Single-cell traces reveal transcriptional bursts. (A) Measured single-cell *her1*-MS2 traces (green) and predicted Her1 protein levels (blue) using the production/decay model from Figure 1C (Methods). Examples of multiple bursts giving rise to a single predicted protein oscillation are highlighted with black arrows. Shaded error bars reflect uncertainty due to background fluctuations and spot detection (Methods). (B) Zoom in to one burst in the last trace of (A). (C) Single z-slices images showing examples of the rapid disappearance and reappearance of a *her1*-MS2 spot, corresponding to the burst highlighted in (B). The z-slice of minute 59 is the same as the previous time point. Spots are highlighted with white arrowheads. Green=her1-MS2; MCP-mNG. Magenta=h2b-mScarlet. (D) Example of promoter state inference via trace binarization. (E) Distribution of number of bursts per predicted protein oscillation for two embryos. 8 ± 3% of oscillations have multiple bursts within them. (F) Distribution of oscillation periods for *her1*-MS2 traces (green) and the predicted Her1 protein signal (blue). Protein oscillations are predicted to be more regular than the measured transcriptional oscillations as revealed by the more peaked distribution of protein periods.

These sharp, noisy pulses are reminiscent of transcriptional bursts in other unicellular and multicellular systems (Lammers, Kim, et al. 2020; Tantale et al. 2016; Chubb et al. 2006; Muramoto et al. 2012; Fukaya, Lim, and Levine 2016; Lee, Shin, and Kimble 2019). Canonical bursts are stochastic (Lammers et al. 2020), at odds with the notion of regular oscillations. However, unlike canonical bursty systems, the segmentation clock is strongly regulated by auto-repression: Her1 proteins repress *her1* transcription (Fig. 1B). This feedback loop clearly could produce or modulate bursts in such a way as to generate regular oscillations. To shed light on the mechanism by which Her1 regulates its own transcriptional bursting and how stochastic transcriptional bursts might interplay with auto-repression to generate noisy but regular oscillations, we turned to mathematical modeling.

### Burst timing statistics offer a window into molecular mechanisms of repression

We extended the classic two-state bursting model to include feedback that accounts for the auto-repression of *her1* (Fig. 6A), ignoring interactions with other clock components such as *her7* (Appendix, Section 3). Feedback can occur via regulation of transcription rate, (‘amplitude regulation’) (Fig. 6B), burst frequency (‘frequency regulation’) (Fig. 6C), burst duration (‘duration regulation’) (Fig. 6D), or a combination thereof. We found that quasi-regular oscillations can emerge from the sole regulation of either amplitude frequency, or duration (Fig. S6). This observation points to the possibility of dynamically rich regulatory strategies, where multiple aspects of bursts are modulated to achieve oscillations, and calls for theoretical predictions that can be tested to rule in or out each strategy.

**Figure 6:**
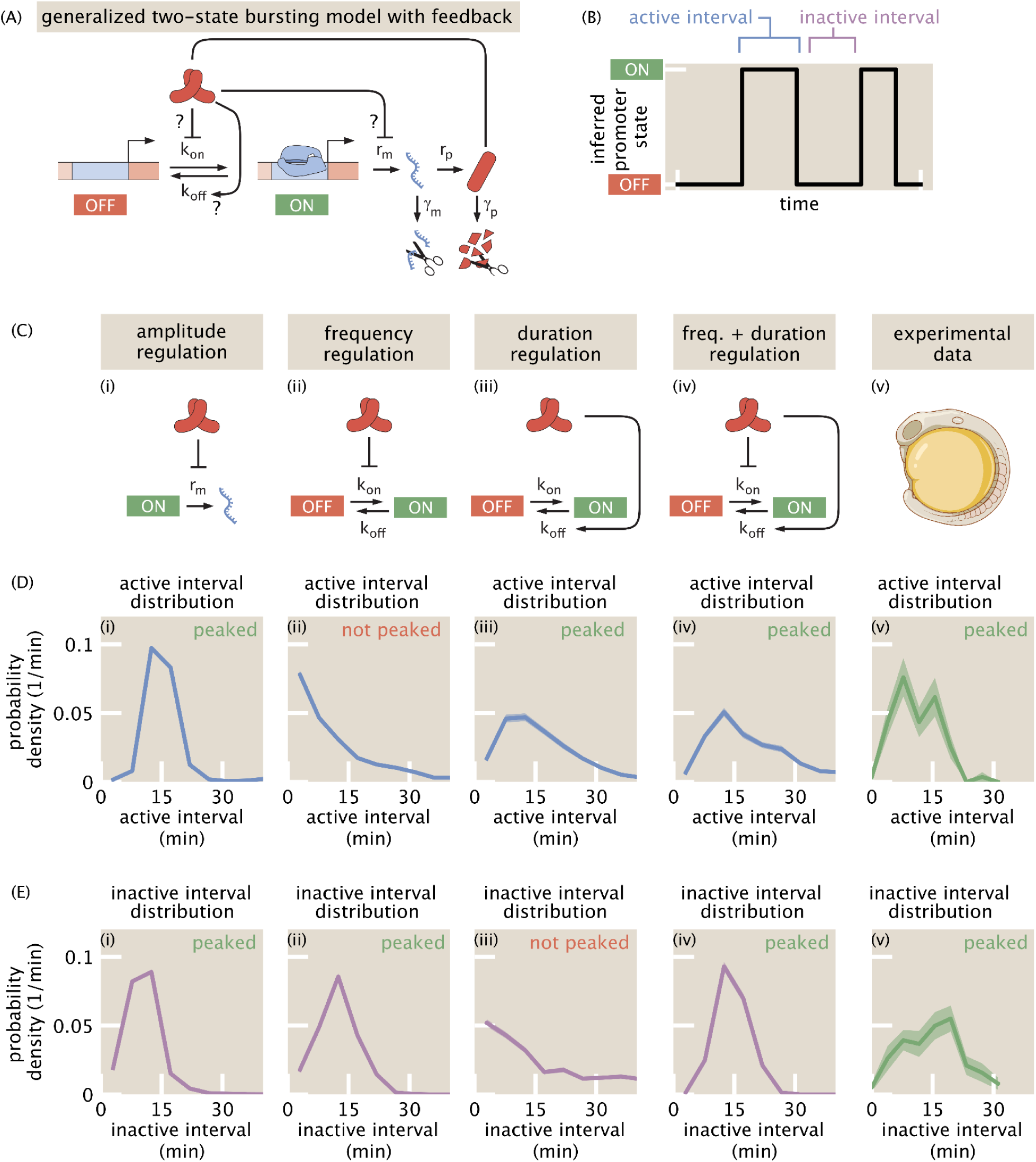
A two-state model of transcriptional bursting with feedback suggests multiple mechanisms of repression could generate oscillations. (A) Schematic of the bursting model. The promoter switches between active (ON) and inactive (OFF) states with rates *k_on_* and *k_off_*, respectively. In the active state, mRNA is transcribed at rate *r_m_*and then translated into protein. Schematic of the active and inactive intervals of bursts. (C) The protein can modulate any of *r_m_, k_on_*, and *k_off_* to repress mRNA production and generate oscillations, which we term amplitude (i), frequency (ii), and duration (iii) regulation respectively. We also consider the case of combined frequency and duration regulation (iv). Our goal is to find evidence for or against these mechanisms in our experimental data from the zebrafish embryo (v). (D)-(E) Simulated (i-iv) and measured (v) active (E) and inactive (F) interval distributions, with simulations covering different mechanisms of repression and including a simulated MS2 reporter with measurement noise (see the section “Generalized bursting model with feedback” in the Methods). Only models with amplitude regulation or combined frequency and duration regulation produce peaked distributions for both active and inactive intervals, as seen in the data. When only frequency or duration is regulated, one of the interval distributions becomes exponential, even accounting for a wide range of measurement noise parameters (Fig. S7-10)

As a first step in uncovering the mechanisms regulating *her1* bursts, we sought to identify signatures of the different regulatory strategies of bursting dynamics in the distribution of active and inactive intervals of bursts (Fig. 6E). The active interval, also known as burst duration, is the length of time that the promoter is in the ON state. The inactive interval is the interval over which the promoter is in the OFF state. In an unregulated two-state model, the distributions of both intervals are exponential, reflecting the Poisson-nature of the ON and OFF switching events. At the opposite extreme, regular oscillations have narrow, peaked interval distributions that become more peaked as the oscillations become perfect. We reasoned that when *k_on_* is regulated but not *k_off_*, the active interval (controlled by *k_off_*) will follow an exponential distribution while the inactive interval (controlled by *k_on_*) will be peaked to allow for bursts to occur in a periodic fashion. Alternatively, for *k_off_* regulation, the inactive interval will be exponentially distributed, while the active interval will be peaked.

To assess whether these differences in the shape of burst interval distributions could be used in practice to distinguish between mechanisms of burst regulation, we measured the active and inactive interval distributions in stochastic simulations that included a model of the MS2 measurement process, choosing model parameters based on the literature where possible (see the “Generalized bursting model with feedback” section of the Methods). As expected, models that produce quasi-regular oscillations (Fig. S6) using only frequency or duration regulation have one peaked and one monotonic interval distribution (Fig. 6F-G). These results are robust to a wide range of simulated measurement noise parameters (Fig. S7-10). In contrast, simulated *her1*-MS2 signals from models that combine frequency and duration regulation or have only amplitude regulation produce peaked distributions for both the active and inactive intervals. Note that in the case of amplitude regulation, the auto-repression itself creates effective “ON” and “OFF” promoter states that are measured by MS2 (Fig. S6). Therefore, these differing predictions for burst timing statistics offer an opportunity to uncover which regulatory strategies are at play, even in the presence of realistic measurement noise.

Turning to the data, we found peaked distributions for both active and inactive intervals (Fig. 6F-G). This result rules against a model of purely frequency or duration regulation, but calls for either amplitude regulation, a combination of frequency and duration regulation, or a combination of all three regulatory mechanisms.

## Discussion

In this work, we established the MS2-MCP mRNA labeling system for quantifying transcriptional dynamics in zebrafish embryos, to our knowledge, a first in any intact vertebrate animal. To facilitate reproducible and quantitative measurements of RNA polymerase activity, we generated a stable line of *ubiquitin*-driven MCP-mNeonGreen that remedies aggregation issues seen in previous lines (Campbell et al. 2015) and avoids many of the issues that accompany approaches in which the transgene is transiently injected for each experiment (Roberts et al. 2014). This line will aid the expansion of transcriptional dynamics research throughout the zebrafish community, as it can be combined with any MS2 construct. To robustly extract single-cell transcriptional dynamics across a millimeter of highly motile tissue (the elongating tailbud), we developed a platform of light sheet fluorescence microscopy and custom image analysis routines that can process terabyte-scale datasets in a few hours. We used this technology to measure the dynamics of *her1*, a genetic oscillator in the core of the segmentation clock. In contrast with the smooth, sinusoidal oscillations seen at the protein level (Delaune et al. 2012; Soroldoni et al. 2014), at the transcriptional level oscillations are discrete and burst-like. Due to the low-pass filtering effects of RNA and protein accumulation, these high frequency bursting dynamics get smoothed out at the protein level. Thus, our finding highlights the power of the MS2 system in being able to reveal novel dynamical features of gene regulation that would remain invisible to classic approaches relying on fluorescent protein fusions.

Our observation of segmentation clock oscillations being composed of discrete transcriptional bursts raises the question of how these bursts are created and regulated to ensure proper protein dynamics that will ultimately drive vertebrate segmentation. In a simple mathematical model, we found that quasi-regular oscillations could be produced if Her1 protein modulated burst amplitude, frequency, or duration, or a combination thereof. Studying burst timing distributions provided evidence in favor of either a mechanism of amplitude regulation, or a combination of frequency and duration regulation, and ruled out a model of purely frequency or duration regulation. In future work, we will directly identify the mechanism of regulation by measuring *her1* transcription while knocking down Her1 protein, either through mutants or morpholinos, and looking for changes in burst amplitude, frequency, or duration. Such an experiment is only possible with RNA labeling technologies like MS2: as demonstrated in Figure 1D,E, these different mechanisms can give rise to identical protein dynamics and are only distinguishable at the level of transcriptional dynamics. Our tools thus present a unique opportunity to dissect the transcriptional regulation of an auto-repressive system.

In characterizing the relationship between transcriptional bursts of *her1* and the predicted Her1 protein oscillation, we found examples of multiple bursting events within one protein cycle. This observation challenges the simple picture of *her1* transcription rate being smoothly modulated by approximately sinusoidal Her1 protein oscillations (Lewis 2003), but rather points to the existence of more complicated and faster transcriptional dynamics. However, one potential caveat to this result is the possibility that the 4 copies of the *her1*-MS2 reporter construct inserted during transgenesis are activating independently and asynchronously. However, the fact that the majority of *her1*-MS2 traces are regular, with one transcriptional burst per protein oscillation, suggests that the 4 copies are strongly coupled such that they act largely in unison. This interpretation is consistent with recent work on the pairing of *her1* and the adjacent clock gene *her7*, which showed that when *her1* and *her7* are on the same chromosome (in *cis*) they are more likely to transcribe concurrently than when they are on opposite chromosomes (in *trans*) (Zinani et al. 2020). For future iterations of the *her1*-MS2 reporter, new landing site technologies for the insertion of transgenes in a single copy would make it possible to circumvent any of the potentially confounding effects of multiple copy insertions (Mosimann et al. 2013; Bhatia et al. 2021; Hadzhiev et al. 2016; Lalonde et al. 2024).

Regardless, most of our results are largely insensitive to the reporter copy number. The tissue-scale dynamics of Figure 4C average over individual cells, washing out small-scale fluctuations that could result from asynchronous firing of multiple reporter copies. The distributions of active and inactive transcriptional intervals in Figure 6F,G would be sensitive to different copies of the promoter firing asynchronously if asynchronous bursts were frequent. However, the fraction of multiple burst events per protein oscillation is small (∼8%). Therefore, even if we assume that all of these multiple burst events are the result of the individual and asynchronous firing of multiple copies of the reporter and exclude them, the overall shape of the active and inactive interval distributions would be unaffected. Finally, the measurement most affected by multiple copies would be the distribution of the number of bursts per protein oscillation (Fig. 5E).

One further limitation of our results is that, at present, it is unclear to what extent one can compare the amplitude of our MS2 signals from different nuclei. Systematic error in the overall fluorescence intensity of spots can arise from multiple sources, including cell-cell variability in MCP expression from our ubiquitin promoter, variable light scattering with tissue depth, and non-uniformity of the excitation beam in light sheet imaging. Exploring alternative promoters with less variability for driving MCP expression will be a useful future endeavor. Further, while the quantification of fluorescence with laser-scanning confocal microscopy is relatively well established, future work is needed to more rigorously assess how to maximize the accuracy of in vivo quantitative fluorescence measurements done with light sheet microscopy. Addressing these issues will be important for inferring the degree of burst amplitude regulation present in the segmentation clock, though should not affect inference of burst timing regulation.

Outside the segmentation clock, the role of transcriptional bursting in genetic oscillators is relevant to a wide range of phenomena. Interestingly, we are aware of two other examples of MS2 measurements of oscillating genes, both of which report bursts within protein-level oscillations. Hafner et al. measured transcription in the p53 pathway and found that approximately 30% of p21 oscillations (period ∼5 hours) contained multiple bursts (Hafner et al. 2020). Wildner et al. studied transcription in the yeast gene CUP1 in response to metal stressors and inferred bursting on the timescale of minutes within oscillations of period ∼30 minutes (Wildner et al. 2023). Together with our results, these findings suggest that transcriptional bursting may in fact be a common feature of genetic oscillators.

Beyond zebrafish, we expect several of the challenges faced here to also be present in other vertebrate systems, such as organoids and mouse pre-implantation embryos. The large size and photosensitivity of these systems make light sheet fluorescence microscopy a useful imaging approach, which comes with well-known challenges of image processing and analysis. Further, these tissues are also often dense and possess strong, structured fluorescence backgrounds that complicate particle detection (Fu et al. 2023). Our image analysis pipeline for MS2 spot quantification, along with the plasmids for MCP and MS2 constructs, may be a useful starting point for such endeavors.

In sum, we envision the tools presented here–in combination with theoretical models–will be useful for expanding the field of quantitative developmental biology further into the domain of vertebrate tissues, where they will help us understand how highly dynamic gene expression programs achieve robustness, and how they fail to do so in disease states.

## Supporting information

Zipped Supplemental Data Files

## Acknowledgements

H.G.G. was supported by an NIH R21 Award (R21HD107436) and the Koret-UC Berkeley-Tel Aviv University Initiative in Computational Biology and Bioinformatics. H.G.G. is also a Chan Zuckerberg Biohub Investigator (Biohub – San Francisco). H.G.G. and A.C.O were also supported by the Human Frontiers Science Program (RGP0041). E.E. was supported by an NSF GRFP (DGE 1752814) and UC Berkeley Chancellor’s Fellowship. B.H.S. was supported by a James S. McDonnell Complexity Fellowship. B.M. was supported by a Chan Zuckerberg Biohub Collaborative Postdoctoral Fellowship. L.A.R., J.B., M.L., X.Z., and S.V. were funded by Chan Zuckerberg Biohub – San Francisco (CZB SF). A.C.O., C.L., and G.V. were supported by the Swiss Federal Institute of Technology in Lausanne EPFL. A.C.O. was also supported by the Francis Crick Institute. A.C.O. was supported by the Max-Planck-Gesellschaft. A.C.O. and G.V. were supported by the Wellcome Trust Senior Research Fellowship in Basic Biomedical Science (WT098025MA). Experiments using the Zeiss Z.1 light sheet fluorescence microscope and Zeiss LSM 880 confocal microscope with AiryScan were conducted at the CRL Molecular Imaging Center, RRID:SCR_017852. We thank the fish facilities at UC Berkeley and EPFL, Tyler Mentley for help in maintaining fish lines, and Arianne Berkowsky for discussions on imaging and image processing.

## Methods

### Ethics statement

All experiments with zebrafish were done in accordance with protocols approved by The University of California, Berkeley’s Animal Care and Use Committee and following standard protocols (protocol number 2018-01-10640-1) and at the EPFL fish facility, which has been accredited by the Service de la Consommation et des Affaires Vétérinaires of the canton of Vaud – Switzerland (VD-H23). All zebrafish used in this study were embryos in the somitogenesis stage of development. Sex differentiation occurs later in zebrafish development (Westerfield 2007) and thus was not a factor in our experiments.

### Zebrafish

Transgenic zebrafish lines of *her1*-MS2 and MCP-mNeonGreen (Fig. 2A) were generated using I-SceI meganuclease-mediated transgenesis (Soroldoni, Hogan, and Oates 2009). Briefly, DNA was co-injected with I-SceI meganuclease into the cell of one-cell stage wild-type embryos (AB for *her1*-MS2, TL for MCP-mNeonGreen). Post-injection, I-SceI enzyme activity was assayed by electrophoresis. One day later, embryos were screened to confirm the fluorescence of mKate2 (a marker of the *her1*-MS2 transgene; see below) or mNeonGreen, whichever was appropriate to the injected DNA. Injected embryos were raised to adulthood and outcrossed to wild-type (AB or TL) zebrafish. If offspring exhibited fluorescence, they were selected to be raised for experiments and their parent was selected as a founder.

The MCP construct uses a ubiquitin promoter (Mosimann et al. 2011) and encodes a ten amino acid linker between the coat protein and fluorescent protein.

The MS2 reporter transgene uses the shared regulatory region between *her1* and *her7*, along with the *her1* coding sequence (Soroldoni et al. 2014). The *her7* coding sequence was replaced with nuclearly localized mKate2, a red fluorescent protein, which serves as a marker of transgenesis (and the presence of MS2) during embryo screening. The *her1* promoter drives the expression of the *her1* coding sequence followed by *MS2v5* (Wu et al. 2015) inserted just before the *her1* 3’UTR.

Sequences for all constructs used in this work can be found at https://benchling.com/garcialab/f_/f9Cist7M-zebrafish-ms2-paper/.

Zebrafish lines were created at the Max Planck Institute of Molecular Cell Biology and Genetics in Dresden and then transferred to the UC Berkeley Zebrafish facility and to the EPFL Zebrafish facility, where they were raised with standard procedures.

To incorporate a stronger nuclear marker, we crossed the MCP-mNeonGreen fish with a line carrying h2b-mScarlet (O’Brown, Megason, and Gu 2019). The resulting offspring were screened for the presence of both fluorophores on a Zeiss AxioZoom widefield microscope.

For imaging, the combined MCP-mNeonGreen; h2b-mScarlet fish were crossed with *her1*-MS2 and the resulting embryos were collected in the morning and then transferred to 19°C after they reached the shield stage. The next morning, embryos were screened for mNeonGreen and mScarlet, and transported from UC Berkeley to the CZ Biohub for light sheet imaging. The presence of the *her1*-MS2 reporter was screened visually on the light sheet via observation of spots.

### Measurement of transgene copy number by qPCR

#### DNA extraction

A subset of transgenic embryos was split into 3 groups of 3 embryos each. Wildtype siblings were split in the same way. An additional group of 10 wildtype siblings was also separated to be used as calibration curves for DNA concentration and to assess primer efficiency. Genomic DNA was extracted from each group of embryos using the DNeasy Blood & Tissue Kit (Qiagen) following the manufacturer’s instructions, but resuspending the DNA in 75 ul of milliQ water instead of the suggested 200 ul of TE buffer. An RNase treatment was done during the extraction. Extracted DNA was stored at -25 °C until the moment of use as substrate for quantitative real-time PCR (qPCR).

#### qPCR

Extracted DNA was prepared to be used as template for qPCR amplification using the SYBR Green MasterMix (ThermoFisher) following the manufacturer’s instructions. Primers targeting *her1* and a reference gene (eukaryotic translation elongation factor 1 alpha 1, *ef1a*) were synthetized (Microsynth). The primer sequences were as follows: her1-qPCR1_F : TCG ATT GGA CAC ATG AGA GC, her1-qPCR1_R : GAA TGG AGG AGA GCT GCT TG, ef1a_F : TCC ACC ACC ACC GGC CAT CT, ef1a_R : CGT GCT GCG CCG CCA TTT T. The qPCR was performed in an Applied Biosystems® QuantStudio 7 (ThermoFisher). Each DNA extraction, including the serial dilution of wildtype embryos, was assessed by triplicate. The qPCR program used was: 50°C (2 min), 95°C (10 min) and 36 cycles of 95°C (15 s), 72°C (1 min). In all cases the temperature gradient was 1.6 °C/s.

#### Estimation of the copy number

The copy number of the *her1*-MS2 transgene was calculated using the relative quantification method proposed by Pfaffl (Pfaffl 2001). Primer efficiencies for *her1* (target gene) and *ef1a* (reference gene) were determined from standard curves generated using serial dilutions of DNA extracted from the pool of 10 wildtype embryos. For each biological replicate, the relative gene dosage of the target gene was calculated by doing the ratio of the efficiency-adjusted Ct values of the target and reference genes. These normalized ratios were then averaged across biological replicates for each genotype. The fold change in relative target gene abundance between transgenic and wild-type samples was used to estimate transgene copy number, under the assumption that each gene copy contributes equally to the qPCR signal: Transgene copies = 2 * Fold Change – 2. With this method, the copy number of the transgene *her1*-MS2 was estimated to be 6 ± 1 (mean ± SEM). In all cases, the Ct values were the average of 3 technical replicates.

### Light sheet imaging

Embryos were imaged using OpenSimView, a 4-objective multiview home-made light sheet microscope (Royer et al. 2016). Embryos were dechorionated using sharp forceps, embedded in 0.1% low melting point agarose (dissolved in E3 media), and pipetted inside optically clear FEP tubes (Zeus Inc. custom order, 2 mm inner diameter) following an established protocol (Kaufmann et al. 2012). E3 media was defined as 5mM NaCl, 0.17mM KCl, 0.33mM CaCl2, 0.33mM MgSO4, pH adjusted to 7.3 with Tris-HCl pH 7.5 1M. A plug of 1.5% agarose was inserted at the bottom of the tube to prevent the leakage of the lower-concentrated agarose. The embryo was oriented such that their tailbud pointed toward the wall of the tube. Once mounted on the microscope, the samples were excited with two light sheets using 488 nm and 560 nm lasers sequentially.

Initial assessment of our ability to detect *her1*-MS2 fluorescent spots was done using a Zeiss Z.1 Light Sheet Fluorescence Microscope (Supplementary Figure 2A), which demonstrated that spots can be readily detected on a standard commercial light sheet fluorescence microscope. However, for time lapse imaging we used our custom 4-objective microscope, which produced higher signal:noise ratios and also was the source of the training data for our nuclear tracking algorithm, which resulted in more accurate tracking.

#### Multiview light sheet details

The light sheets were generated using two Nikon N10XW-PF objectives (0.3 NA, 3.5 mm WD). Stacks of 400 z-slices with a spacing of 0.97 μm and a frame interval of 1 minute were acquired during 3 hours at 28 C. The total size of the field of view was 388×1000×1000 μm (*z*-*y*-*x*), with a pixel size of 0.485 μm in *x-y*. Images were acquired using two Nikon N16XLWD-PF detection objectives (0.8 NA, 3.0 mm WD) and two Hamamatsu ORCA-Flash 4.0 V3 Digital CMOS cameras. The use of two lightsheets and two cameras results in four images per channel for each time point, which are later computationally fused as described below.

During image acquisition, we encountered a hardware communication issue that led to a blocked filter wheel and subsequently missed images for approximately 10% of time points. We removed these blank time points but accounted for the missing time intervals. The rest of the scans were unaffected.

## Data and code availability

Code for the full image analysis pipeline can be found at https://github.com/GarciaLab/zms2.

The output of this code for the two datasets in this paper along with two manually curated datasets and additional files is included as Supplementary Data Files.

Code for reproducing the plots in this paper, including model simulations, can be found at https://github.com/GarciaLab/zebrafish-ms2-paper.

### Image processing

The raw images were first converted to the *zarr* format (Miles et al. 2020). This format divides the image data into smaller tiles (i.e., chunks) that are compressed on the disk and are loaded on demand, allowing faster data loading and avoiding excessive memory compared to TIFF. The *zarr* format also addresses the problem of concurrent data access present in the *h5* file format (Miles et al. 2020). Then, for each time point, the four images that constitute each pair of light sheets and cameras were fused using the *dexp* Python package (Yang et al. 2022) as done in (Lange et al. 2024). In brief, the fusion is done in two steps. First, the volumes from different light sheets and the same camera are fused, resulting in one volume per camera, two volumes in total. These two volumes are registered to compensate for a minimal misalignment between the cameras and then fused, resulting in a single volume. During fusion, small differences in the overall brightnesses of the two camera views are corrected for by linearly rescaling the dimmer image to best match the 1% and 99.9% quantiles of the brighter image. After fusing the images, the intensity of the Histone channel was normalized across the time series. A Gaussian blur and background subtraction were also applied during this step to facilitate nuclear segmentation, but no further processing was done to the MCP channel.

### Nuclear segmentation and tracking

Cell nuclei and their boundaries were detected using a 3D U-NET (Ronneberger, Fischer, and Brox 2015; Jordao Bragantini et al. 2024). Detected nuclei were segmented and tracked over time using the approach described in (Jordão Bragantini, Lange, and Royer 2023; Jordao Bragantini et al. 2024) using the following parameters: *threshold*=0.25, *max_area*=7500, *min_area*=200, and *min_frontier*=0.0. Both the segmentation and tracking were performed on a desktop computer (Intel Core i9 11900K, GeForce RTX 3090 24GB, 128 GB RAM) running Ubuntu 20.04.

### Spot analysis

To perform spot analysis, we developed a custom Python pipeline. Our pipeline takes in two *zarr* arrays, one corresponding to the MCP channel images and the other corresponding to a label matrix that encodes the nuclear segmentation and tracking information, and outputs a *pandas* DataFrame with spot location and intensity information.

The goal of the spot analysis is to locate all of the true MS2 spots and quantify their intensity. The pipeline is organized into the following steps (Supplemental Figure 3A):

1. *Detection:* a first pass of detecting spots is performed using simple filtering and thresholding.
2. *Classification:* voxels containing detected spot-like objects are classified into ‘spot’ and ‘not spot’ using a convolutional neural network classifier which outputs a probability of belonging to the ‘spot’ class. The user provides a threshold probability value to reject false spots (we used 0.7).
3. *Quantification:* the intensity of the remaining spots is measured by estimating a background level of pixels just outside the spot, subtracting this background from spot pixels, and summing the resulting pixel values.

### Detection

First, bright skin cells that surround the presomitic mesoderm were removed from the image using blurring and thresholding. Then, potential spots were identified with a simple Difference of Gaussians filter and thresholding the filtered image, with parameters chosen by visual inspection of a single time point. The filtering and binary thresholding is done on each 2D z-slice independently on a GPU using the *cucim* library. The resulting 3D mask was then transferred back to the CPU for the label matrix calculation. The centroid of each object in the label matrix was computed as an approximate location of the potential spots in the image.

### Classification

A small convolutional neural network was trained to classify voxels as containing a spot or no spot. Voxels were 9×11×11 pixels^3^. The network was implemented in *Keras*. The network architecture was inspired by a model used to classify fluorescence microscopy images of bacteria (Hay and Parthasarathy 2018). Schematically, the network consists of 2 convolution layers followed by a fully connected layer that outputs the probability of containing a spot. Specifically, the first 3D convolutional layer has 4 kernels, each of size 3×3×3 pixels^3^, ReLU activation and ‘same’ padding, followed by max pooling. The second 3D convolutional layer has the same parameters as the first and is also followed by max pooling. We then flatten into a dense layer with 512 neurons and ReLU activation. During training, this layer is followed by a Dropout layer with dropout = 0.5. The result is then fed into a single sigmoid neuron for output.

The small size of this network allows it to be trained on a small number of hand-labeled spots. We manually classified 1250 voxels from 4 time points, resulting in a training dataset with 227 spots and 1023 “not spots”. We trained the model by minimizing the binary cross-entropy loss using the Adam optimizer with a learning rate of 10^-4^ and batch size of 8 in 5-fold cross validation. We augmented the training data with random rotations. To offset class imbalance during training, we used class weights equal to half the inverse fraction of each class in the training data. After 100 training epochs, the validation loss decayed to 0.21 ± 0.05 (mean ± standard deviation) and remained stable (Supplemental Fig 3C). The final average validation area under the curve (AUC) was 0.97 ± 0.01.

During spot analysis, the initial set of detected candidate spots is filtered down using a probability threshold of 0.7.

### Quantification

Spot quantification occurs through particle localization, background estimation and subtraction, and then integration over a defined ellipsoid of size (4×4×4) pixels^3^, or (2×2×4) microns^3^. Our pipeline contains two particle localization algorithms: least-squares Gaussian fitting and the radial center algorithm (Parthasarathy 2012), which is an analytic estimate of particle center based on radial symmetry. Both methods are highly accurate, approaching the Cramer-Rao bound (Parthasarathy 2012). The radial center algorithm is inherently much faster, as it requires no fitting. However, we implemented multi-processor parallelization of *scipy’s least_squares* function that is quite fast in practice. As discussed more below, Gaussian fitting has the benefit that the best-fit width parameters have strong predictive power in further discriminating between true and false spots. Throughout this paper, we used the Gaussian fitting method.

Importantly, because of the background MCP-mNeonGreen expression in and around each nucleus has spatial structure, straightforward application of both particle localization algorithms, which assume uniform background with Poisson noise, fail on the majority of spots. To minimize the structured background, we first perform Difference of Gaussians filtering on the spot voxel and then estimate particle centers.

Next, we estimate the background levels for each spot individually and subtract that background from the spot pixel intensities. Background estimation can be done in one of two ways. The first way is to re-fit a 3D Gaussian to the spot in real space, but with fixed center and width parameters, extracting just an offset and amplitude. Gaussian fitting is the standard way to estimate background levels of MS2 spots (Garcia et al. 2013). The second way is to take a “shell” of pixels of a fixed width and distance from the spot center and average the intensity of the shell. Using a shell of inner width = 4 pixels and outer width = 6 pixels, we find that the two methods of background estimation strongly agree with one another (Supplementary Fig 3F, R^2^ = 0.94). In all data in this paper, we used the shell method, as it is faster.

With the background estimate in hand, we subtract this background from all pixels in the voxel, enforcing non-negativity. We then define an ellipsoid of size 4×4×4 pixels^3^ and sum the background-subtracted intensities.

To assess the accuracy of this quantification procedure, we generated a library of synthetic spots with known integrated intensity and realistic background and noise (Supplemental Fig. 4A). Specifically, we modeled spots using a Gaussian point spread function (PSF) and a constant offset. We then modeled the structured background by adding to this Gaussian an exponential function with decay length equal to 10 pixels and a random orientation in 3D. Finally, we modeled shot noise by assigning to each pixel value a Poisson random number with mean equal to the intensity of the noise-less pixel.

We ran our spot quantification pipeline on this library and assessed intensity accuracy as a function of spot amplitude (which sets the signal-to-noise ratio) and the amplitude of the exponential gradient. We measured the relative accuracy of the intensity measurement, defined as (measured intensity - true intensity) / true intensity, across spot parameters and also using two different spot localization methods: first, Gaussian fitting on the raw pixels, and second, Gaussian fitting on Difference-of-Gaussians (DoG)-filtered images, the latter to remove the structured background. We found that localization using DoG-filtered images led to considerably enhanced accuracy across spot parameters (Supplemental Fig. 4B).

To make contact with the experimental data, we defined two readily measured parameters: signal:background ratio (where signal is the mean background-subtracted spot pixel intensity) and the “structuredness” of the background, defined as the variance-to-mean ratio of the of the shell pixels that define the background. For a uniform background with Poisson noise, the structuredness is 1. Binning our simulations in this 2D space, we created an accuracy regime diagram (Supplemental Fig. 4C). Averaging across all spots in our dataset, we found that the experimental data had a signal:background value of 0.35 ± 0.24 (mean ± standard deviation), i.e., 35% above background, and a structuredness value of 40 ± 21, reflecting the highly structured MCP background visible in the spot images. Nevertheless, with DoG-based localization and our Gaussian combined with our exponential model of spots, we infer that our spot intensity quantification is highly accurate, with an relative error of at most 40%, but typically around 5%.

### Trace assembly

MS2 traces are assembled by assigning spots to nuclei and linking them through time using the nuclear tracking information. First, we check if a spot’s centroid falls within a region identified as a nucleus by the “Segments” label matrix that is output from the nuclear tracking pipeline. If a spot does not fall within a nucleus, then we perform a search for the nearest nucleus within a cube of size 11 pixels in x and y and 7 pixels in z, centered on the spot. If multiple spots are assigned to the same nucleus, we pick the one with the highest probability value from the neural network classifier.

After assigning each spot to a nucleus, we assemble a draft of the traces and begin iteratively refining them. In each iteration we perform the following steps. We identify regions of transcriptional activity by thresholding a moving average of the trace using a window of 5 time points and a threshold of 1 a.u. (effectively any non-zero point passes the threshold). For each point in the “on” region of the trace such that the raw trace has a value of zero and has an adjacent point with a value greater than zero, we attempt to fill in that point. We extract a voxel centered on the location of the adjacent point but at the “empty” time point. We then run this voxel through our spot classification and quantification routines. We accept or reject the spot using either the spot classifier or quantification parameters. For all data in this paper, we kept only spots for which all Gaussian width parameters (sigma_x, sigma_y, and sigma_z) fell within the range 0.5 and 3.0 pixels, which we found to be an effective filter by manual inspection.

Upon iterating this process, the set of found points that pass the given criteria will converge. For all data in this paper, we used 10 iterations.

After the traces are assembled, we can make further cuts by rejecting spots based on spatial location. Previous measurements of protein levels (Delaune et al. 2012) and smFISH counts (Keskin et al. 2018) indicate that *her1* transcription halts after somite boundary formation. For reasons unknown, we found that background levels of MCP-mNeonGreen increase in somites after boundary formation, leading to increased false positive detections in this region. We therefore reject spots that are greater than 40 μm anterior of the last formed somite, as measured along our defined anterior-posterior axis (see below). The position of the last formed somite was determined manually over time. Similarly, we restrict our analysis to spots detected within the PSM and tailbud tissues by rejecting spots that lie greater than 50 μm from the anterior-posterior axis (see below), corresponding to, for example, false detections in the skin. Finally, for analysis of single-cell traces, we keep only traces with greater than 10 spots.

### Spot intensity uncertainty

The uncertainty in individual spot intensity was estimated using a method based on (Garcia et al. 2013). A full derivation of the uncertainty is given in Section 2 of the Appendix. In brief, the uncertainty was assumed to be dominated by the error in background estimation. This error is measured by considering the time evolution of the background around each spot, fitting a mean trend using a 4th order polynomial, and computing the RMS deviation of background levels from this trend. In contrast to the original method (Garcia et al. 2013), we use a 4th order polynomial as opposed to a spline as we found that additional degrees of freedom in the trend model led to worse performance in a cross-validation scheme (Appendix, Fig. A2).

In addition, and in contrast to previous works, due to the increased difficulty of the spot detection problem we further incorporate uncertainty in spot detection via the empirically measured false-positive and false-negative rates (Appendix, Section 2.2). Combining both of these sources of uncertainty, we arrive at the final uncertainty for an MS2 traces as

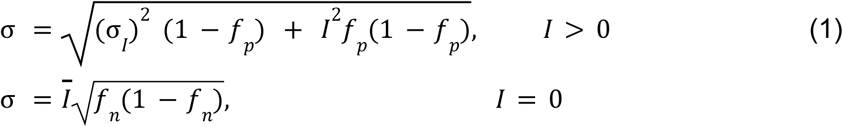

where σ_*I*_ is the uncertainty from background fluctuations, *f*_*p*_ is the false-positive rate of detection, *f*_*n*_ is the false-negative rate of detection, *I* is the intensity of the individual spot, and *Ī* is the mean intensity of the trace. From analysis of manually-curated traces, we measured *f* = 0. 04 and *f* = 0. 08. σ depends on the trace, but is typically 1-10% of *Ī*. See Section 2 of the Appendix for more details.

### Anterior-posterior axis

We define the anterior-posterior axis using a combination of manually placed points and spline interpolation. Using the interactive data viewer *napari* (Sofroniew et al. 2022), we construct a coarse anterior-posterior axis by manually placing around 10 points that span from the last formed somite (which becomes the origin) to the tip of the tail bud in the first time point of the timeseries. We place these points in 3D using a *napari* Points layer, viewing one slice at a time, roughly following the notochord. In the tailbud, we complete the anterior-posterior axis by projecting points towards the closest outer boundary of the tissue. Keeping track of the location of that last formed somite, we proceed to define coarse anterior-posterior axes every 50 time points until the end of the movie. We then refine these anterior-posterior axes through spline interpolation. First, we interpolate the anterior-posterior axis of each time point, going from 10 to 100 spatial positions, using a B spline of degree 2. Then, for each 100 points along the new anterior-posterior axis, we interpolate over time, using a B spline of degree 1. We use *scipy’s splprep* function to define the spline. Spots are assigned to an anterior-posterior axis bin by finding the axis point that has the smallest euclidean distance to the spot’s centroid.

### Protein signal prediction

We use a linear model of transcription, translation, and protein maturation to predict what mRNA (*m*) and protein signal (*p*) would be created by our MS2 traces (*x*), assuming that the fluorescent intensity of the MS2 signal is proportional to the instantaneous rate of transcription (Lammers, Galstyan, et al. 2020). Specifically, we model the system according to

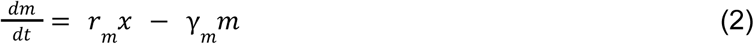

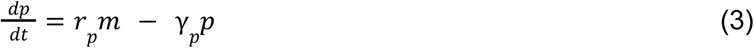

When no spot is detected, the MS2 trace *x* is assigned a value of zero. We numerically integrate this model using *scipy*’s *solve_ivp* function with the RK45 routine and interpolating the MS2 signal, *x*, between observation times using *numpy*’s *interp* function.

The key parameters are the decay rates (*γ_m_*, *γ_p_*) of *her1* mRNA and Her1 protein, respectively, which we take from reference (Lewis 2003) to both be 0.23 min^-1^. The production rates (*r_m_*, *r_p_*) simply set the concentration scales of mRNA and protein, respectively, which here can be taken as arbitrary since we measure mRNA levels and predict protein levels in arbitrary units. The initial conditions of total mRNA and protein are unknown parameters. In Section 1 of the Appendix, we show that the predicted mRNA and protein signals become independent of the initial condition on the time scale of the decay constants. Therefore, for simplicity, we pick initial conditions of mRNA(t=0)=protein(t=0)=0 and ignore approximately the first cycle of predicted signals.

In Fig. 1C-E, we use this model to demonstrate the low-pass filtering effect of the central dogma (more details in Appendix, Section 1). We generated toy example MS2 traces consisting of either a sine wave or a square wave and used these as inputs to the protein prediction model, which outputs smooth oscillations in both cases.

### Kymograph creation and tissue-scale period measurement

To create a kymograph, spots were grouped by their anterior-posterior position into 100 equally spaced bins (approximately 5 μm long). The intensities were summed within each bin and plotted as a heatmap with time increasing downwards. The lengthening of the tissue-scale period was measured directly from the kymograph by summing the intensities from bins located between 240 and 315 μm along the anterior-posterior axis and plotting the resulting signal over time. Oscillation peaks were identified by fitting a Gaussian profile to manually-defined sections of the signal that isolated single oscillations and taking the fit center. Periods were calculated as the differences between successive peaks. Uncertainty in the period was obtained by bootstrapping over spatial bins within the 240-315 μm range. The predicted protein measurements were done identically, but instead of the spot intensities using the simulated protein signals evaluated at the observation time points.

### Burst calling and calculating number of bursts per oscillation

We implemented a conservative burst calling algorithm based on thresholding a moving average of the *her1*-MS2 traces. The moving average was computed via convolution with a uniform kernel of size 3 time points, and we used a threshold of 1 a.u., effectively grouping any non-zero signal into a burst. With this algorithm, fluctuations in *her1* transcription are determined as corresponding to distinct bursts only if they are separated by 3 time points of inactivity. We defined oscillations using the predicted protein signal. We used *scipy’s find_peaks* function to identify peaks in the predicted protein oscillation. We then collected burst start and end times and assigned the bursts starting between the previous protein peak and the current protein peak to the current protein peak. With all bursts being assigned to a protein peak, we then computed a histogram of the number of bursts per protein peak, which we call the number of bursts per oscillation. With our conservative burst calling algorithm, we likely underestimate the frequency of stochastic bursting within individual Her1 protein oscillations.

### Generalized bursting model with feedback

Details of the mathematical model and its implementation are found in the Appendix, Section 3. In brief, we extended the canonical two-state bursting model to allow for protein concentration to increase the rate of transcription, decrease the rate of OFF→ ON events, or increase the rate of ON → OFF events (Fig. 6A) according to Hill functions. The model contains three chemical species: the *her1* gene, which can be in the inactive state, *G*, or the inactive state, *G\**; *her1* mRNA, denoted by *M*; and Her1 protein, *P*, whose total number is denoted by *p*. These three species are governed by the following set of reactions:

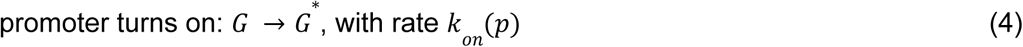

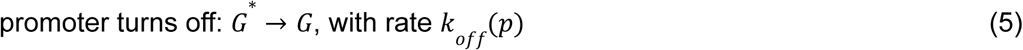

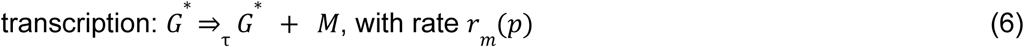

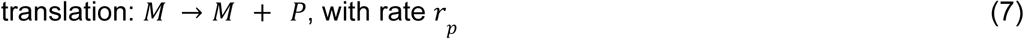

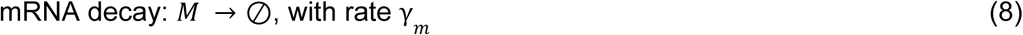

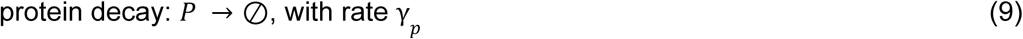

We performed fully stochastic Gillespie simulations of the model, with the mRNA production reaction occurring with a time delay, τ, (denoted by the double arrow with a τ subscript) following (Bratsun et al. 2005).

#### Model parameters

The following parameters were held fixed: mRNA decay rate (*γ_m_*) = 0.23 1/min, protein decay rate (*γ_p_*) = 0.23 1/min (Lewis 2003), maximum transcription rate (*r_m_*) = 10 mRNAs/min (chosen to obtain an oscillation amplitude of approximately 40 mRNAs (Keskin et al. 2018), translation rate (*r_p_*) = 4.5 proteins/mRNA/min. The remaining model parameters were chosen manually to produce simulated MS2 traces that qualitatively resemble the experimental data and exhibit statistically increased regularity in the resulting oscillations compared to unregulated bursting. For the simulations in Figure 6, the following parameters were used:

- (B) Amplitude regulation: the transcription rate is repressed by the protein according to a Hill function, *r_m_*/(1 + (*p*/*K_D_*)*^n^)*. We ignore promoter switching (i.e., the promoter is always in the ON state by setting *k_off_*=0 1/min), letting the sharp repression of the transcription rate itself create burst-like behavior. Parameters: *K_D_*=100 proteins, *n*=3, τ=7.5 min.
- (C) Frequency regulation: *k_on_* is repressed according to a Hill function, *k_on_* = *k^+^*/(1 + (*p*/*K_D_*)*^n^)*, with a maximum value of *k^+^.* Parameters: *k^+^*=0.5 1/min, *k_off_*=0.08 1/min, *K_D_*=80 proteins, *n*=3, τ=0 min.
- (D) Duration regulation: *k_off_* increases with protein level according to an increasing Hill function, *k_off_*(*p*) = *k^-^*(*p*/*K_D_*)*^n^*/(1 + (*p*/*K_D_*)*^n^*), with a maximum value of *k^-^*. Parameters: *k_on_*=0.055 1/min, *k^-^*=0.4 1/min, *K_D_*=1100 proteins, *n*=3, τ=0 min.
- Frequency and duration regulation: both *k_on_* and *k_off_* are Hill functions of protein concentration, with maximum values *k^+^* and *k^-^*, respectively. Parameters: *k^+^*=0.5 1/min, *k^-^*= 0.4 1/min, *K_D,on_*=80 proteins, *K_D,off_*=1100 proteins, n=3, τ=0 min.

#### Simulated MS2 signals

From each simulation we extracted a time series of the promoter state variable, which fluctuates between 0 (“OFF”) and 1 (“ON”). Using these promoter state traces, we simulated realistic MS2 traces using the model of (Lammers, Galstyan, et al. 2020) that encapsulates the effect of RNA polymerases being loaded on and off the gene with a memory kernel of normalized length *w* corresponding to the dwell time of RNA polymerase on the gene as it engages in transcription. Based on the measured elongation rate for *her1* via smFISH (Hanish) 4.8 kb/min and our approximately 1.4kb MS2 cassette at the 3’ end of the transgene, we used a dwell time = 0.29 min, which, with our sampling rate of 1.0 1/min implies w = 0.29. We further included multiplicative Gaussian noise with standard deviation σ*_MS2_*to simulate fluorophore emission and measurement uncertainty, and a detection threshold, below which the simulated MS2 signal was set to zero. We used default values of σ*_MS2_* = 0.2, and detection threshold = 0.2 * *w* * max(*r_m_*), where *r_m_*is the mRNA production rate in equation 6 and is time-dependent in the case of amplitude regulation. These values were chosen because they generated traces with a scale of fluctuations that visually matched our experimental data. However, we also varied these parameters in our robustness analysis (Fig. S7-S10).

#### Protein period regularity measurement

To assess the regularity of simulated protein oscillations in our model, we used the *scipy* find_peaks function with prominence = 0.01 to identify the locations of peaks in protein oscillations. We then defined the protein period as the intervals between peaks and computed the coefficient of variation (CV, standard deviation/mean) of these periods.

## Figures

**Supplemental Figure 1:**
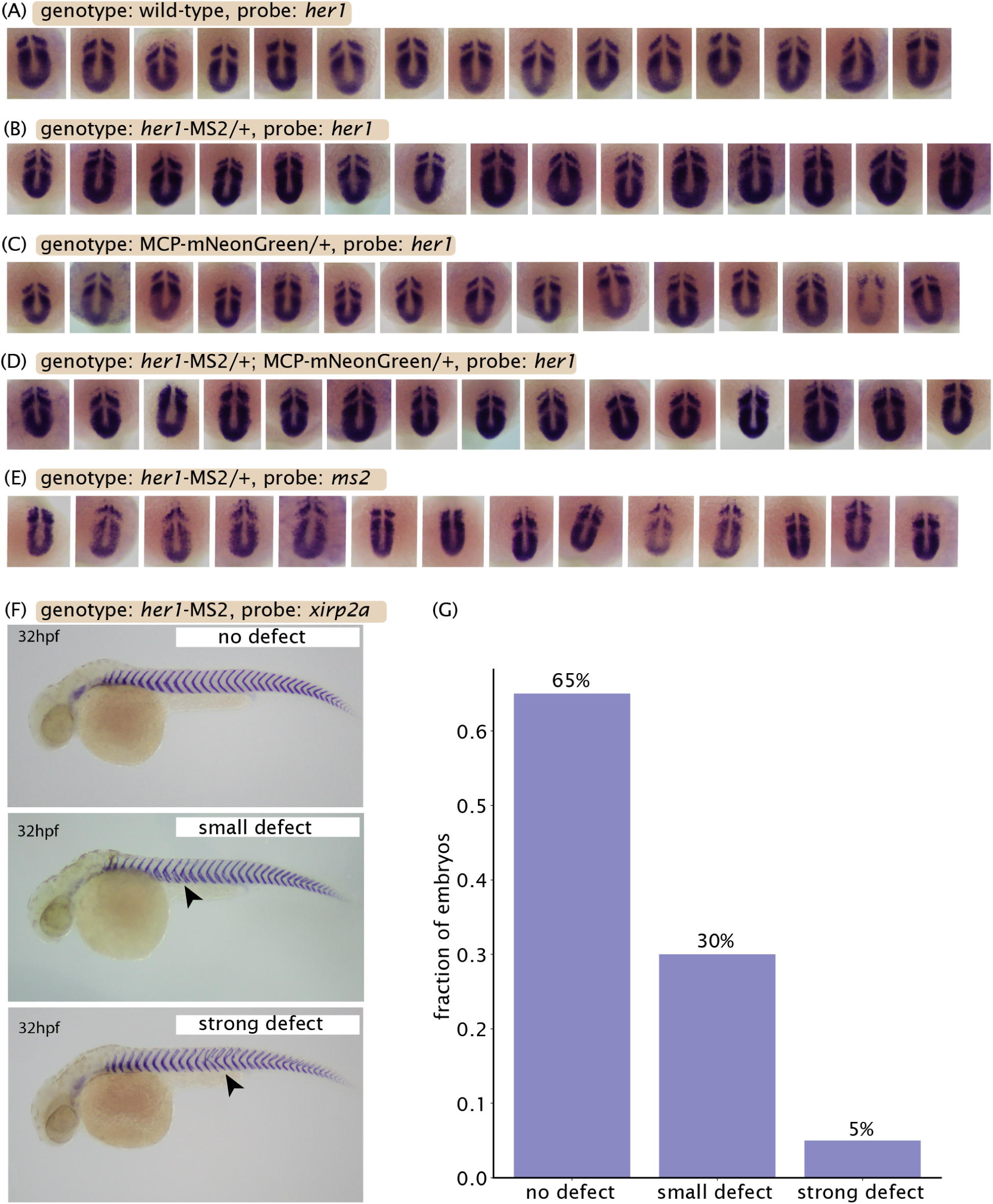
Controls show overall healthy somitogenesis of *her1*-MS2 and MCP-mNeonGreen transgenic lines. (A)-(D): In situ hybridization assays against *her1* and MS2 in various genotypes at the 12-14 somite stage embryos showing qualitative agreement of the *her1* expression pattern in both wild-type and transgenic fish. All embryos are offspring of a cross of *her1*-MS2 and MCP-mNeonGreen heterozygous animals. (A) “Wild-type” embryos lacking both markers for MS2 and MCP show normal *her1* expression patterns (Schröter et al. 2012). (B) “*her1*-MS2” embryos carrying only the MS2 transgene show healthy *her1* expression. “MCP-mNeonGreen” embryos carrying only the MCP transgene show healthy *her1* expression. (D) Embryos carrying both transgenes show healthy *her1* expression. (E) In situ hybridization assay against the MS2 sequence in *her1*-MS2 embryos showing that the reporter qualitatively recapitulates the endogenous *her1* expression pattern. (F) *In situ* hybridization assay for the somite boundary marker *xirp2a* (Riedel-Kruse, Müller, and Oates 2007), showing examples of no defect, small defect, strong defect. When defects do occur, they are limited to the trunk region, which completes somite formation before our imaging experiments begin and is thus not included in our analysis. (G) The majority of animals show no defects at 32 hours post fertilization.

**Supplemental Figure 2:**
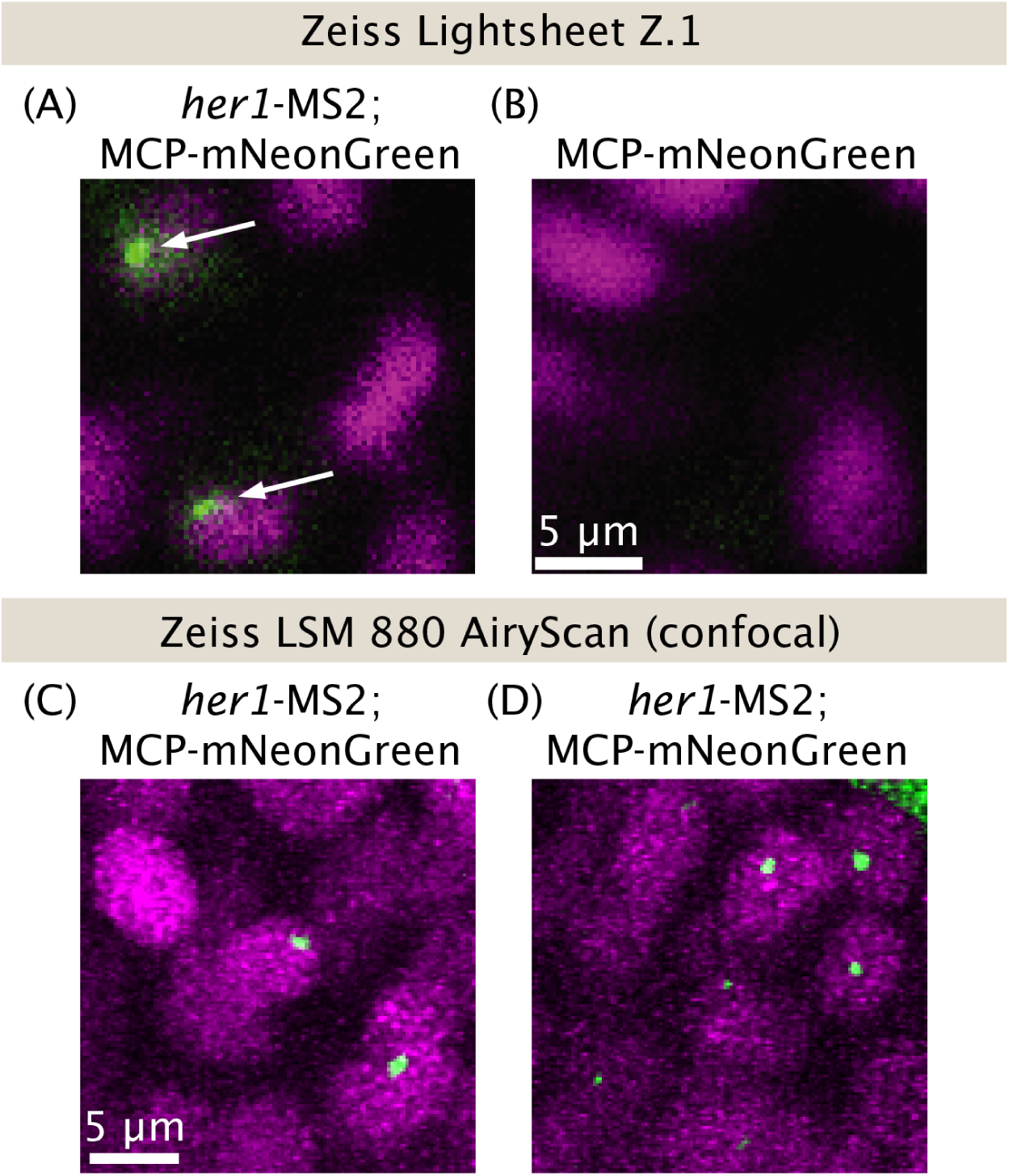
Characterization of spots on commercial light sheet and confocal fluorescence microscopes. (A,B) Spots (white arrows) are detected on the Zeiss Z.1 Lightsheet but only when the reporter gene is present. MCP-mNeonGreen fish show no puncta in the absence of the *her1*-MS2 reporter. Single z-slices of embryos with MCP-mNeonGreen (green) and h2B-mScarlet (magenta) as a nuclear marker (A) with *her1*-MS2 or (B) without *her1*-MS2. Contrast was adjusted manually such that background MCP levels were approximately equal in both images. (C)-(D) Two examples of spots on a Zeiss LSM 880 confocal microscope with AiryScan processing.

**Supplemental Figure 3:**
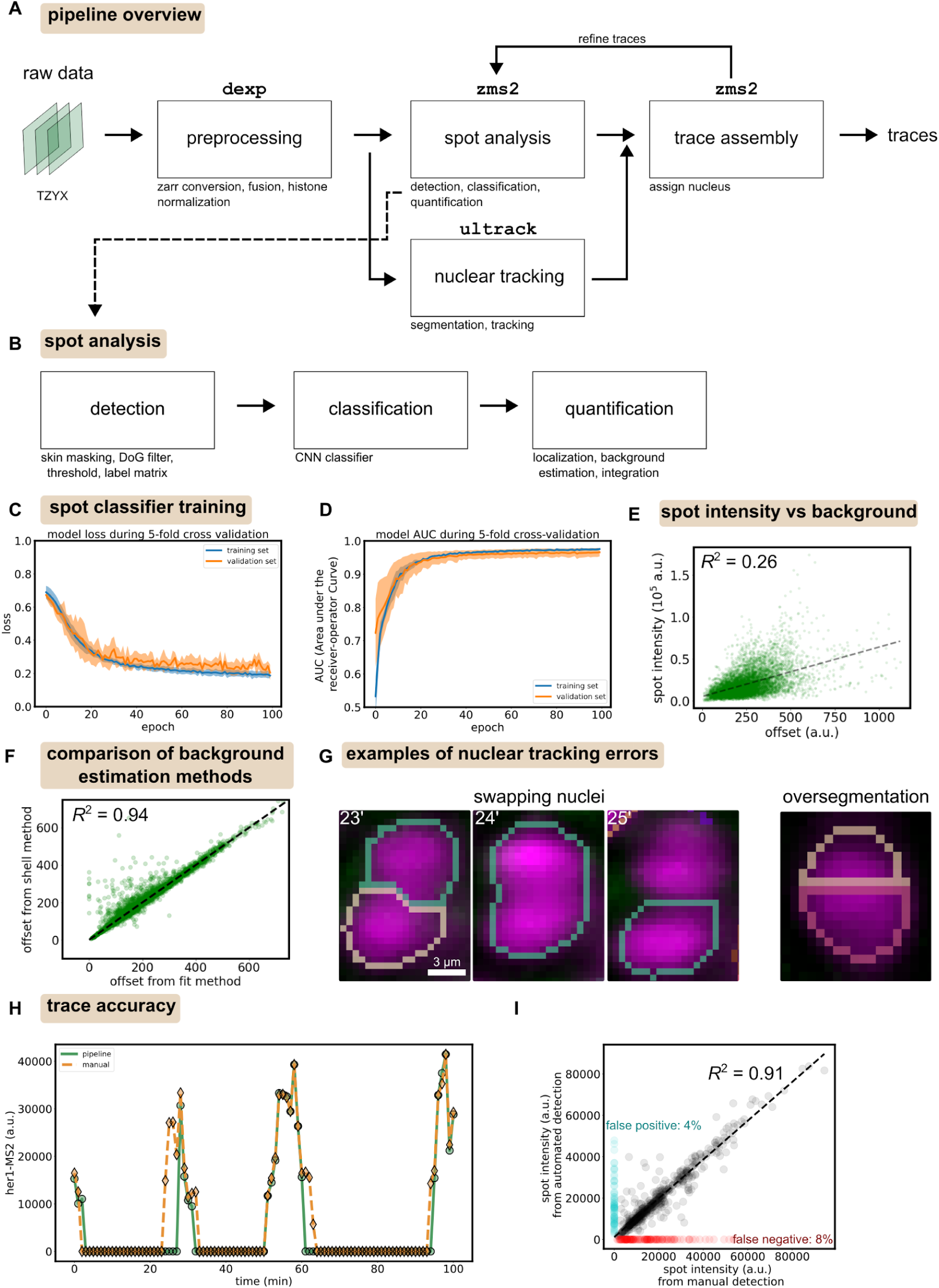
Overview of image analysis pipeline and its accuracy. (A) Schematic of the full pipeline. The pipeline takes two 4D image stacks as input, one for the nuclear marker and one for the MS2-MCP reporter per light sheet view. Preprocessing using the *dexp* package (Yang et al. 2022) involves conversion to the *zarr* file format, fusion of multiple light sheet views (if necessary), and normalization of the nuclear marker channel to prepare for nuclear segmentation. After preprocessing, the nuclear and spot channels are split, with the nuclear channel undergoing segmentation and tracking via *ultrack* (Jordao Bragantini et al. 2024) and the spot channel passing through our spot pipeline, called *zms2*, which detects and analyzes MS2 spots. The outputs of these packages are combined into traces of spot fluorescence for a given nucleus as a function of time. Traces are then refined using an iterative computational algorithm. In brief, the algorithm identifies abrupt gaps in traces and attempts to fill them in by extracting a voxel around the expected spot location, passing it back through the classification and quantification modules, and seeing if the spot passes a set of criteria (see the “Spot analysis” section of the Methods). (B) Schematic spot analysis section of the full pipeline. Candidate spots are detected and then classified as a true spot or a false-positive detection using a binary Convolutional Neural Network (CNN) classifier. The culled list of spots then undergo quantification, where the total fluorescence intensity of the signal is computed via spot localization, background estimation and subtraction, and then summing pixel values within a defined ellipsoid (see the “Spot analysis” section of the Methods). DoG=Difference of Gaussians, CNN=Convolutional Neural Network. Skin masking=computational removal of bright and irrelevant skin cells. Thresholding occurs on the DoG-filtered image to create a binary mask, which is transformed to a label matrix that assigns a unique identifier to each candidate spot. (C) Binary cross-entropy loss function of convolutional neural network spot classifier over training shows that the model converged with minimal overfitting. Solid lines are means and shaded error bars standard deviations over 5 folds of cross validation. Blue=training set (“train”), Orange=validation set (“val”). CV=cross validation. (D) Area Under the receiver-operator Curve (AUC), a metric of model performance that balances false positive and false negative detections, during training (“epoch” = time in the training process). The validation AUC converged to 0.97 ± 0.01, indicating very high classification performance (a perfect classifier has AUC=1, an unbiased random guess has AUC=0.5). (E) Spot intensity vs. background level (“offset”) showing an extremely weak correlation (*R^2^*=0.26), which is indicative of MCP saturation and reliable measures of absolute fluorescence intensity. (F) Comparison of two methods for background estimation, described in the Methods section. “Shell” refers to taking the mean of pixel values in a shell around the spot. “Fit” refers to fitting a Gaussian with an offset. We find that the two methods give very similar results, with the data clustering closely around the 1:1 line (black dashed line) and an *R^2^* of 0.94. The best fit slope is 1.049 +/- 0.003. (G) Examples of nuclear tracking errors, including when the algorithm swaps the identities of nearby nuclei (left), and when it splits a single nucleus into two segments (right). Shown are single z-slices of histones (magenta), with the algorithm-assigned identity of each nucleus marked by a unique colored outline. MCP-mNeonGreen background is faintly visible in green. (H) Example comparison of a pipeline-derived trace (green solid line) and the corresponding manually-derived trace (orange dashed line), obtained by manually clicking on spots in images and passing these spots through the quantification pipeline. (I) Comparison of spot intensities between pipeline and manual traces across 728 spots from 26 nuclei. Cases where both methods detect a spot are colored in black. Linear regression to these points shows strong correlation (R^2^ = 0.91). Cases in which a human detected a spot and the pipeline did not (false negative) are colored in red. These cases are 8% of all spots. Cases in which a human did not detect a spot but the pipeline did (false positive) are colored in cyan. These cases are 4% of all spots.

**Supplemental Figure 4:**
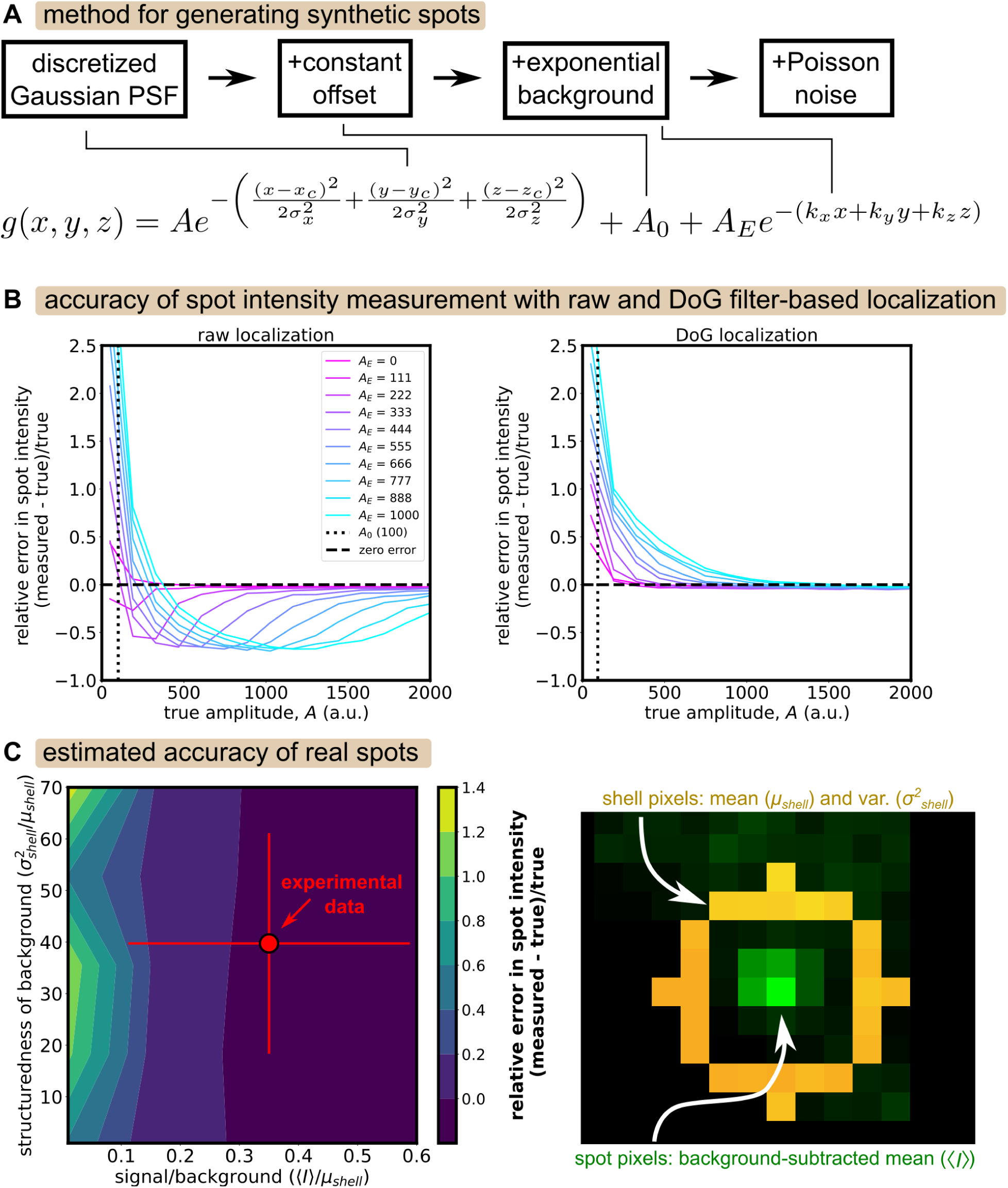
Characterization of spot intensity measurement accuracy using synthetic spots. (A) Schematic of method for generating synthetic spots with structured background. The Point Spread Function (PSF) is modeled as a Gaussian function, to which we add a constant offset and an exponentially decaying background, the latter being a simple model of the structured MCP background we observe in real spots (not shown). The wavevector, *k*, of the background is taken to have a length of 10 pixels (roughly the length of the spot in voxels) and a random direction. The pixels of the final image are then drawn from a Poisson distribution with mean equal to *g(x,y,z)*. (B) Relative error in spot intensity (measured - true)/true vs true spot amplitude (the parameter *A* in the equation in panel (A)) for different strengths of the exponential background (via the parameter *A_E_*). Left: spot localization via Gaussian fitting of the raw image. Right: spot localization via Gaussian fitting of a Difference-of-Gaussians (DoG)-filtered image. After localization, background estimation is done with the “shell” method described in the Methods. Intensity is then the sum of background-subtracted pixels inside a defined ellipsoid. Intensity errors are larger when localization is done on the raw pixels, due to the structured background. (C) Left: Placing the experimental data (red circle=mean, red bars=standard deviation across all spots) in a regime diagram describing relative accuracy of the intensity measurement (heatmap) as a function of signal/background and “structuredness” of the background. The x-axis, signal/background, is the average background-subtracted value of spot pixel intensities, e.g., 0.1 = 10% above background. The y-axis, structuredness of background, is the variance of shell pixel intensity divided by the mean of the shell pixel intensity . For a uniform background with Poisson noise, the structuredness will be 1. The experimental data has a background with structuredness of approximately 40 on average. The simulations from (B) were binned into a 2D histogram according to their signal/background and structuredness levels to create the final heatmap of relative error in measured spot intensity, defined as (measured intensity - true intensity)/(true intensity). For example, a relative error of 1.0 means the algorithm overestimated the total spot intensity by 100%, or a factor of 2. This heatmap illustrates that, based on our model of spots, the DoG-based localization method should lead to highly accurate intensity estimations, with a relative error of less than 10%. Right: Schematic of the parameters used to define the axes of the heatmap on the left, using a single z slice of a real spot (green) with shell pixels highlighted in orange.

**Supplemental Figure 5.**
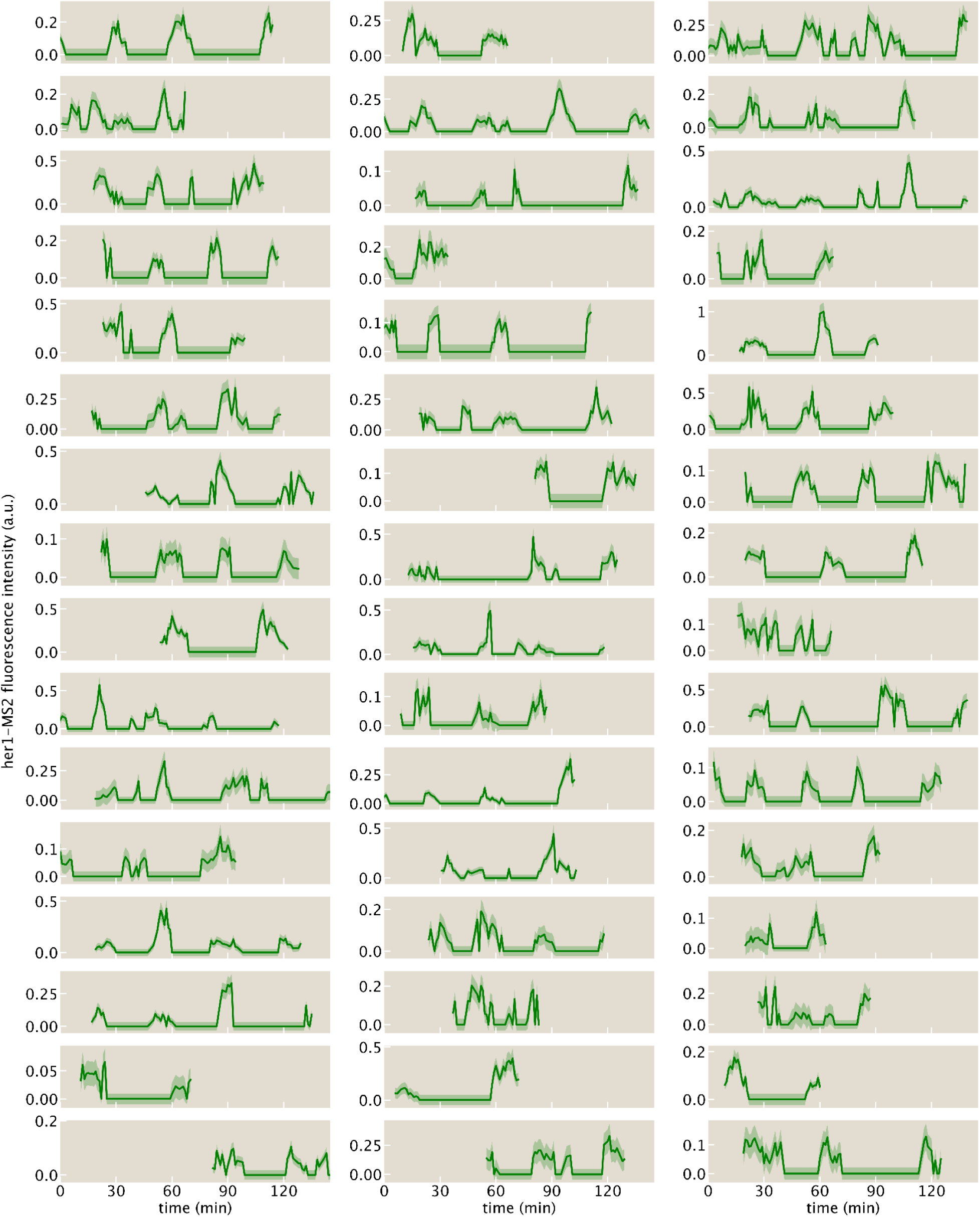
48 randomly chosen traces with at least 20 spots per trace.

**Supplemental Figure 6.**
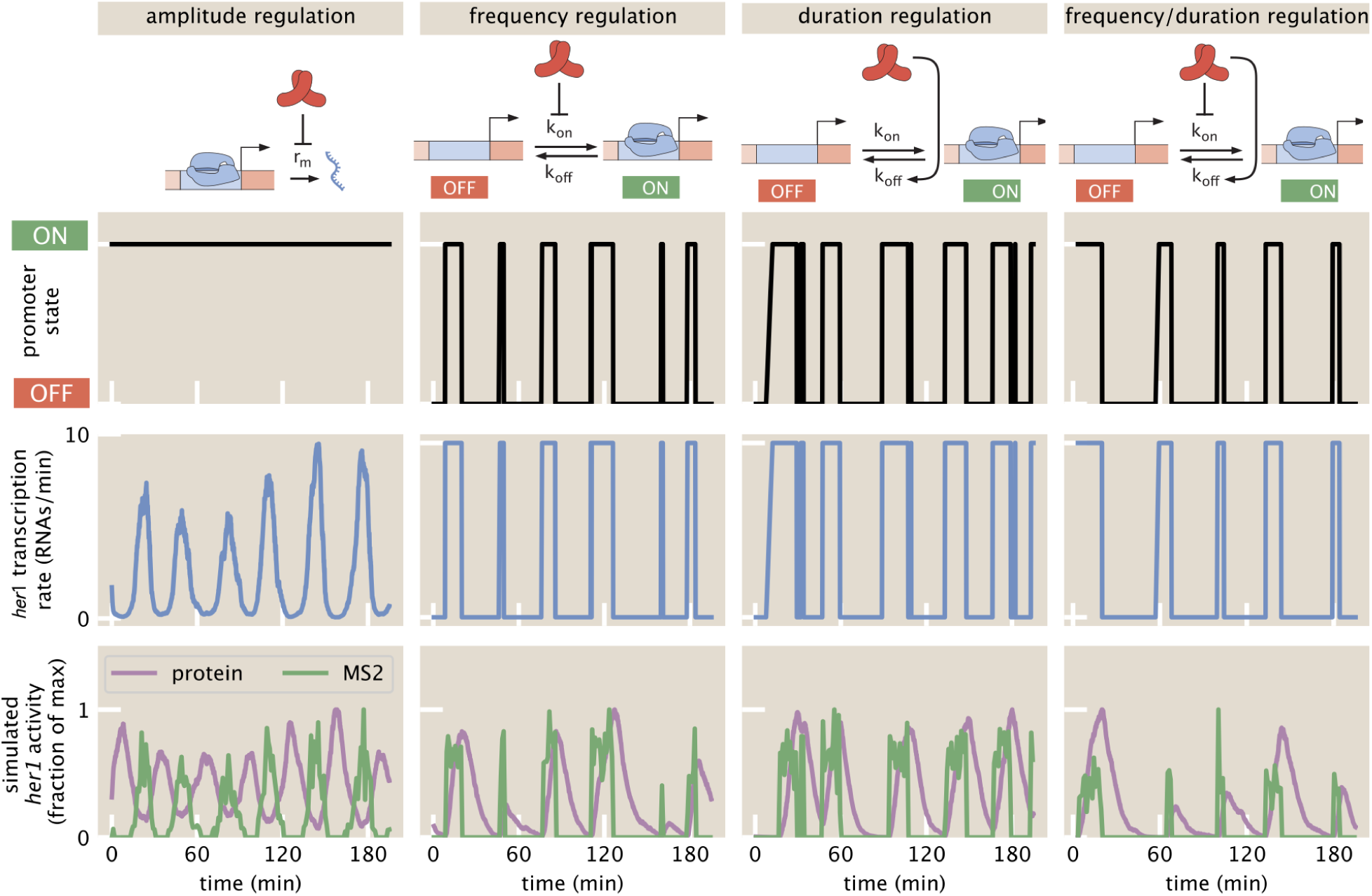
Examples of simulated traces for different models. Each column corresponds to a different model of autorepression (see Fig. 6). Top row: dynamics of promoter state switching. Middle row: instantaneous transcription rate. Bottom row: simulated traces of the expected *her1*-MS2 signal (Methods) and Her1 protein concentration, normalized to maximum. Parameters are the same as in Fig. 6, described in the Methods section.

**Supplemental Figure 7:**
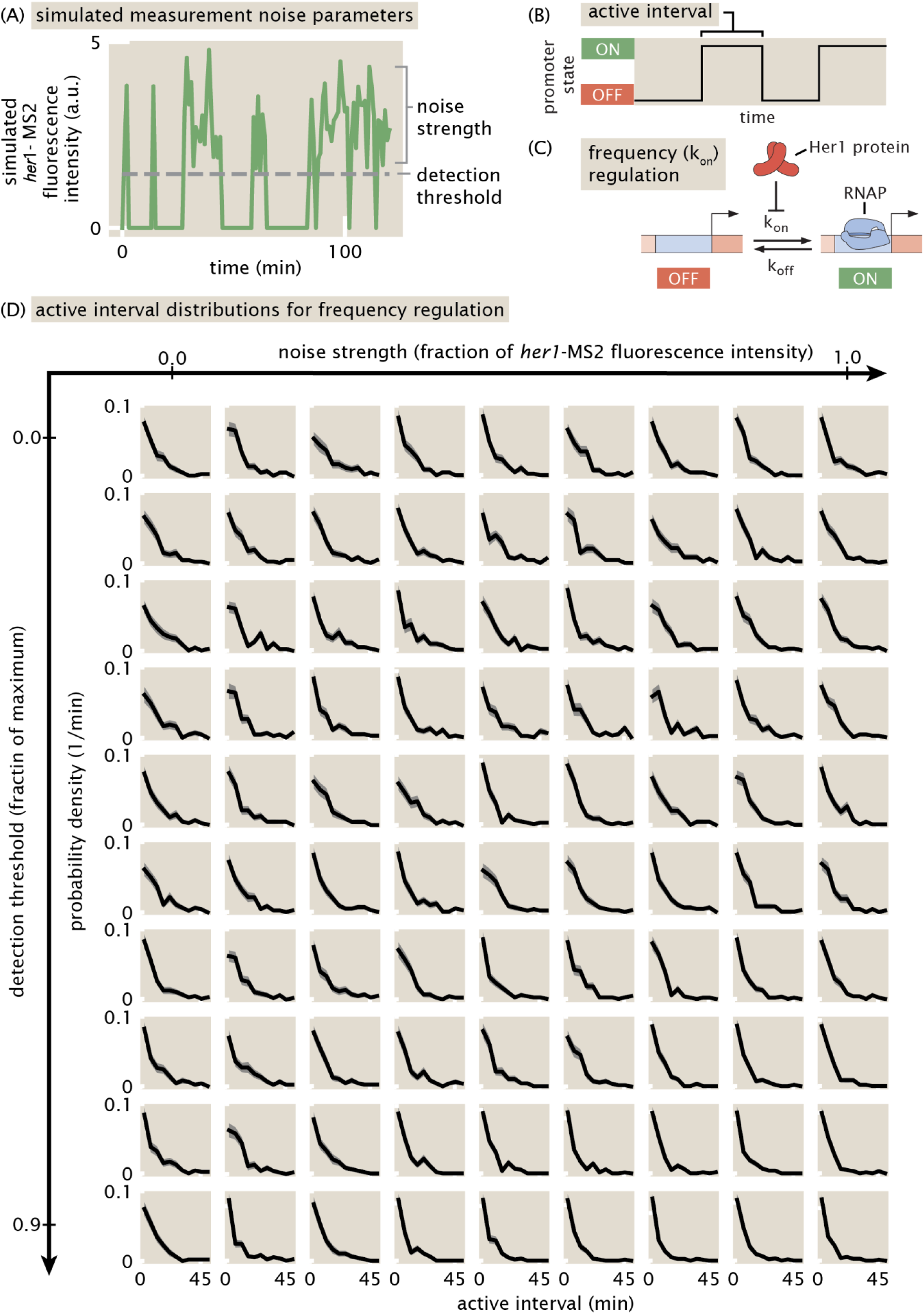
Active interval distributions for the frequency regulation bursting model remain exponential for a wide range of simulated measurement noise parameters. Our goal is to assess the impact of the MS2 measurement process on our ability to correctly identify the shape of burst interval distributions. (A) Simulated *her1*-MS2 trace illustrating the two measurement noise parameters: noise strength, which determines the level of measurement noise within a burst, and detection threshold, which determines how strong the *her1*-MS2 signal must be to be detected. Specifically, “noise strength” is the amplitude of multiplicative Gaussian noise added to the simulated *her1*-MS2 trace; we vary the noise strength from 0.0 to 1.0. The “detection threshold” is implemented by setting all values less than the detection threshold to zero; we vary the detection threshold from 0.0 to 0.9 times the maximum *her1*-MS2 signal. Other model parameters are described in the “Generalized bursting model with feedback” section of the Methods. These parameter values were chosen to produce quasi-regular oscillations while being consistent with the literature where possible. (B) In this figure, we focus on the analysis of the active burst interval (also known as burst duration), defined as the duration of time the promoter is in the ON state. (C) We also focus here on the case of frequency regulation, where Her1 proteins regulate the rate of the promoter turning on, *k_on_*. In the absence of measurement noise, the active interval distribution for frequency regulation is exponential, since the active interval is set by *k_off_*, not *k_on_*, and here *k_off_* is constant. (D) We explored how measurement artifacts affect the shape of this distribution by creating a 2D grid of active interval distributions, with increasing noise strength to the right and increasing detection threshold going down. The active interval distribution remains exponential over all of the explored parameter space, indicating that the degree of measurement noise explored here does not change the distribution’s qualitative shape.

**Supplemental Figure 8:**
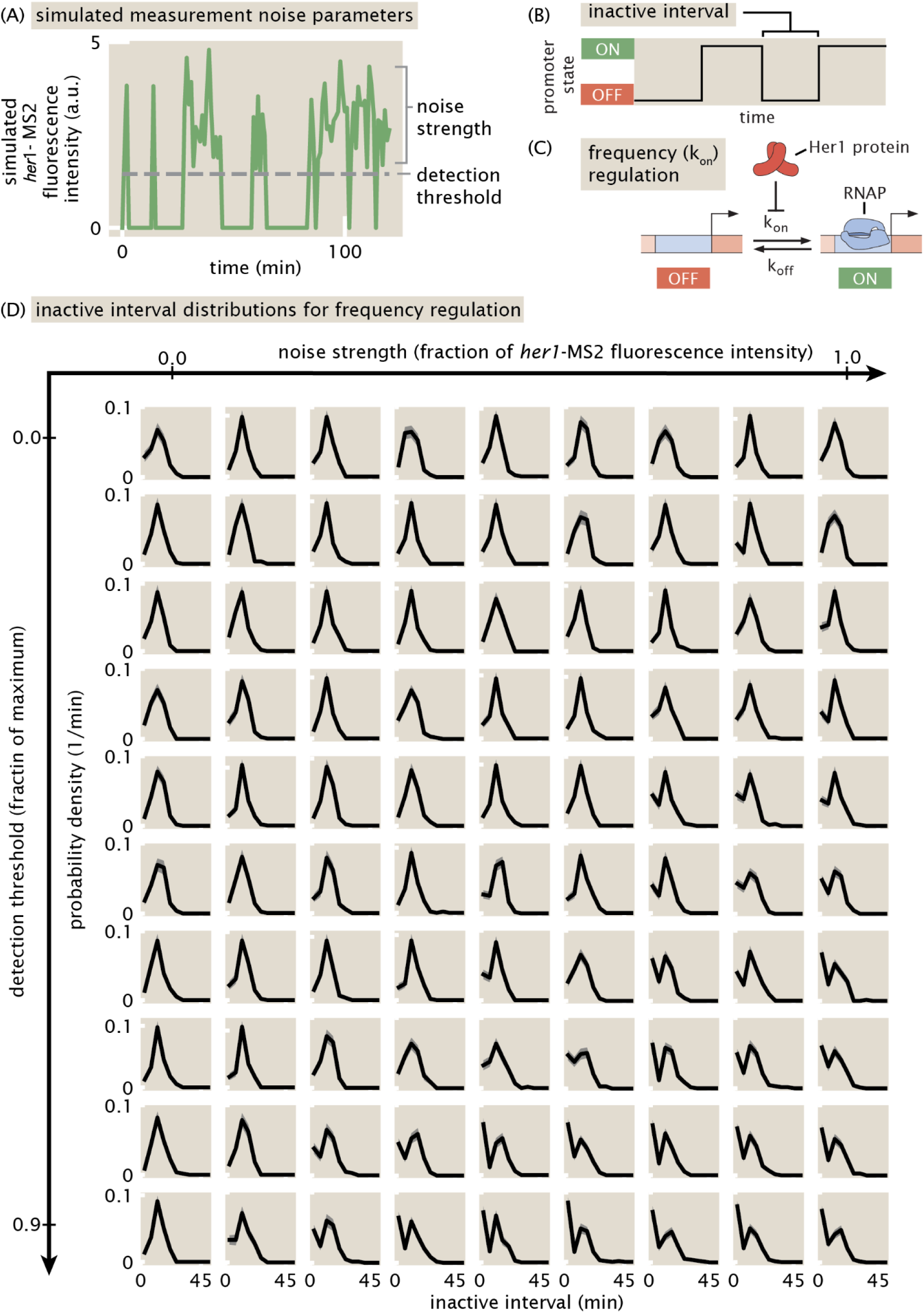
Inactive interval distributions for the frequency regulation bursting model remain peaked for a wide range of simulated measurement noise parameters. Our goal is to assess the impact of the MS2 measurement process on our ability to correctly identify the shape of burst interval distributions. (A) Simulated *her1*-MS2 trace illustrating the two measurement noise parameters: noise strength, which determines the level of measurement noise within a burst, and detection threshold, which determines how strong the *her1*-MS2 signal must be to be detected. Specifically, “noise strength” is the amplitude of multiplicative Gaussian noise added to the simulated *her1*-MS2 trace; we vary the noise strength from 0.0 to 1.0. The “detection threshold” is implemented by setting all values less than the detection threshold to zero; we vary the detection threshold from 0.0 to 0.9 times the maximum *her1*-MS2 signal. Other model parameters are described in the “Generalized bursting model with feedback” section of the Methods. These parameter values were chosen to produce quasi-regular oscillations while being consistent with the literature where possible. (B) In this figure, we focus on the analysis of the inactive burst interval, defined as the duration of time the promoter is in the OFF state. (C) We also focus here on the case of frequency regulation, where Her1 proteins regulate the rate of the promoter turning off, *k_off_*. With our chosen parameters and in the absence of measurement noise, the inactive interval distribution is peaked, reflecting the regulation of *k_on_* that reduces stochasticity in burst initiation. (D) We explored how measurement artifacts affect the shape of this distribution by creating a 2D grid of inactive interval distributions, with increasing noise strength to the right and increasing detection threshold going down. The distribution remains peaked over a wide range of the explored parameter space, with high noise strength and/or high detection threshold adding an exponential background to the distribution.

**Supplemental Figure 9:**
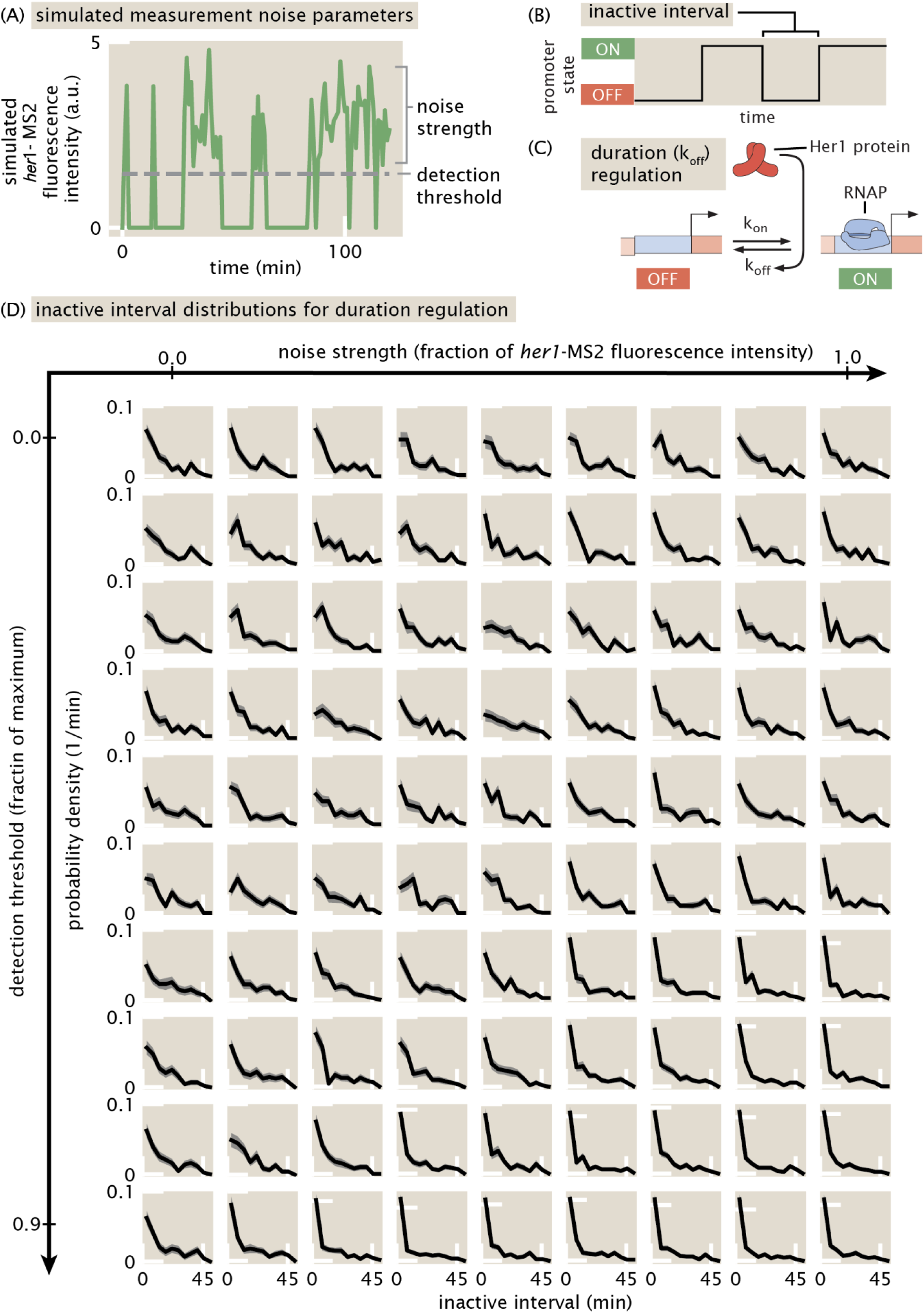
Inactive interval distributions for the duration regulation bursting model remain exponential for a wide range of simulated measurement noise parameters. Our goal is to assess the impact of the MS2 measurement process on our ability to correctly identify the shape of burst interval distributions. (A) Simulated *her1*-MS2 trace illustrating the two measurement noise parameters: noise strength, which determines the level of measurement noise within a burst, and detection threshold, which determines how strong the *her1*-MS2 signal must be to be detected. Specifically, “noise strength” is the amplitude of multiplicative Gaussian noise added to the simulated *her1*-MS2 trace; we vary the noise strength from 0.0 to 1.0. The “detection threshold” is implemented by setting all values less than the detection threshold to zero; we vary the detection threshold from 0.0 to 0.9 times the maximum *her1*-MS2 signal. Other model parameters are described in the “Generalized bursting model with feedback” section of the Methods. These parameter values were chosen to produce quasi-regular oscillations while being consistent with the literature where possible. (B) In this figure, we focus on the analysis of the inactive burst interval, defined as the duration of time the promoter is in the OFF state. (C) We also focus here on the case of duration regulation, where Her1 proteins regulate the rate of the promoter turning off, *k_off_*. In the absence of measurement noise, the inactive interval distribution for duration regulation is exponential, since the inactive interval is set by *k_on_*, which is constant here. (D) We explored how measurement artifacts affect the shape of this distribution by creating a 2D grid of inactive interval distributions, with increasing noise strength to the right and increasing detection threshold going down. The distribution remains exponential over all of the explored parameter space.

**Supplemental Figure 10:**
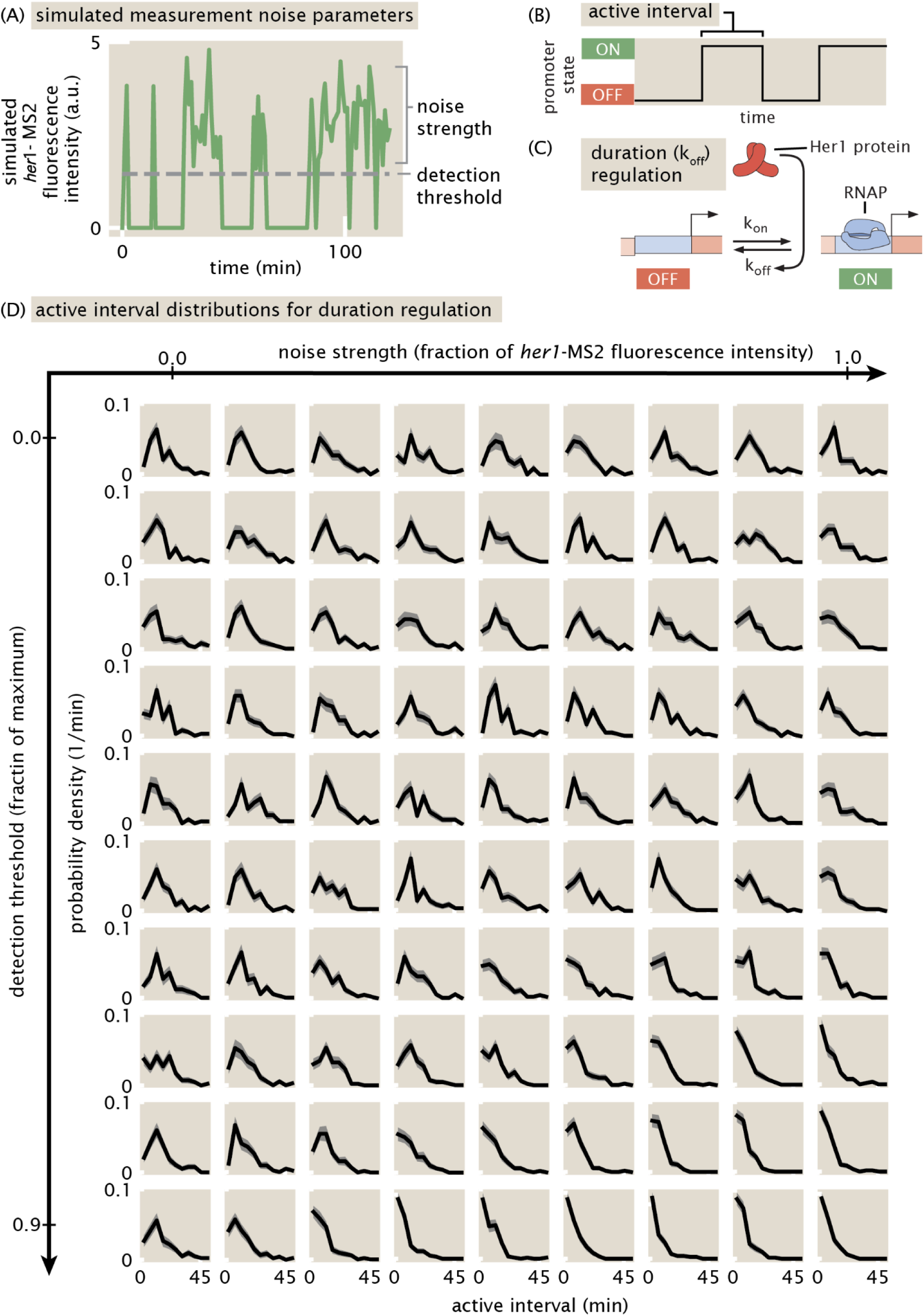
Active interval distributions for the duration regulation bursting model remain peaked for a wide range of simulated measurement noise parameters. (A) Our goal is to assess the impact of the MS2 measurement process on our ability to correctly identify the shape of burst interval distributions. (A) Simulated *her1*-MS2 trace illustrating the two measurement noise parameters: noise strength, which determines the level of measurement noise within a burst, and detection threshold, which determines how strong the *her1*-MS2 signal must be to be detected. Specifically, “noise strength” is the amplitude of multiplicative Gaussian noise added to the simulated *her1*-MS2 trace; we vary the noise strength from 0.0 to 1.0. The “detection threshold” is implemented by setting all values less than the detection threshold to zero; we vary the detection threshold from 0.0 to 0.9 times the maximum *her1*-MS2 signal. Other model parameters are described in the “Generalized bursting model with feedback” section of the Methods. These parameter values were chosen to produce quasi-regular oscillations while being consistent with the literature where possible. (B) In this figure, we focus on the analysis of the active burst interval (also known as burst duration), defined as the duration of time the promoter is in the ON state. (C) We also focus here on the case of duration regulation, where Her1 proteins regulate the rate of the promoter turning off, *k_off_*. With our chosen parameters and in the absence of measurement noise, the active interval distribution for duration regulation is peaked, reflecting the regulation of *k_off_* that reduces stochasticity in burst duration. (D) We explored how measurement artifacts affect the shape of this distribution by creating a 2D grid of active interval distributions, with increasing noise strength to the right and increasing detection threshold going down. The distribution remains peaked over a wide range of the explored parameter space, with high noise strength and/or high detection threshold adding an exponential background to the distribution that eventually dominates.

## Supplemental Movies

<u>Supplemental Movie 1:</u> Maximum intensity projections of AiryScan confocal microscopy images of zebrafish embryo with nuclei shown in red and *her1*-MS2 in green. Due to cell motion, manual adjustment of the microscope stage was required every few minutes to keep the same cells in the field of view. These manual adjustments correspond to the discrete jumps in the image observed throughout the movie.

<u>Supplemental Movie 2:</u> 3D rendering of light sheet fluorescence microscopy images of a zebrafish embryo with nuclei false colored according to the label assigned to them by the nuclear tracking algorithm. To better resolve somites, skin nuclei were computationally removed. Some flickering in the movie occurs due to imperfect segmentation of the skin. Aside from this surface-level flickering, the overall stability of colors conveys the high level of tracking accuracy.

<u>Supplemental Movie 3:</u> Rotating 3D rendering of light sheet fluorescence microscopy images of a zebrafish embryo with nuclei false colored according to the label assigned to them by the nuclear tracking algorithm. To better resolve somites, skin nuclei were computationally removed. Some flickering in the movie occurs due to imperfect segmentation of the skin. Aside from this surface-level flickering, the overall stability of colors conveys the high level of tracking accuracy.

<u>Supplemental Movie 4:</u> 3D rendering of light sheet fluorescence microscopy images of a zebrafish embryo with nuclei shown in gray and false-colored in proportion to their *her1*-MS2 signal.

<u>Supplemental Movie 5:</u> Rotating 3D rendering of light sheet fluorescence microscopy images of a zebrafish embryo with nuclei shown in gray and false-colored in proportion to their *her1*-MS2 signal.

<u>Supplemental Movie 6:</u> Animated 3D rendering of light sheet fluorescence microscopy images of a zebrafish embryo with nuclei shown in gray and false-colored in proportion to their *her1*-MS2 signal (left) and in proportion to their predicted protein signal (right). See Methods for details of protein prediction.

<u>Supplemental Movie 7:</u> Animated 3D rendering of light sheet fluorescence microscopy images of a zebrafish embryo with nuclei shown in magma and the anterior-posterior axis, defined through a combination of manual labeling and spline interpolation, shown in white spheres.

## Appendix

See Supplemental Appendix File

### Supplemental Data Files

**Supplemental Data File 1: Dataset_1.pkl**. *pandas* DataFrame that is the output of the image analysis pipeline (images from the Dorado light sheet microscope). Each row corresponds to an MS2 spot. Columns contain various spot features. See the code for details. This dataset was used for Figures 3 and 4.

**Supplemental Data File 2: Dataset_1_Curated.pkl**. *pandas* DataFrame that is the output of the image analysis pipeline (from the Dorado light sheet microscope) and manually curated, with each spot visually confirmed and linked to the correct nucleus. Each row corresponds to an MS2 spot. Columns contain various spot features. See the code for details. This dataset was used for the single-cell analysis in Figures 5 and 6.

**Supplemental Data File 3: Non_Blank_Time_Points.pkl**. *numpy* array that contains the true scan numbers of each time point in the Dorado dataset. As discussed in the Methods section, hardware communication issues during acquisition led to missing images for ∼10% of scans. These blank time points were removed from the image dataset, but the proper timing information is stored in the non_blank_timepoints array.

**Supplemental Data File 4: Dataset_1_Nuclear_Tracks.csv**. .*csv* file that contains the location and identity of every tracked nucleus.

**Supplemental Data File 5: Simulated_Intervals.pkl**. Pickled lists containing the simulated active and inactive intervals used in Figure 6. The order of the data is: amplitude regulation, frequency regulation, duration regulation, frequency and duration regulation. For each regulation mode there are 3 lists: active intervals, inactive intervals, and periods. See also the corresponding Fig. 6 notebook on the paper’s github repository for the simulation code.

## Notes

### Competing Interest Statement

The authors have declared no competing interest.

### Summary of Updates

Revised analysis of single-cell burst distributions; clarified figures.

