## Appendix for "Single-cell transcriptional dynamics in a living vertebrate"

April 21, 2025

1. Biophysics Graduate Group, University of California at Berkeley, Berkeley, USA
2. Department of Molecular and Cell Biology, University of California, Berkeley, CA, USA
3. Department of Physics, University of California, Berkeley, CA, USA
4. Institute for Quantitative Biosciences-QB3, University of California, Berkeley, CA, USA
5. Chan Zuckerberg Biohub – San Francisco, San Francisco, CA, USA
6. Institute of Bioengineering, EPFL; Lausanne, CH
7. Department of Cell and Developmental Biology, UCL; London, UK
8. The Francis Crick Institute; London, UK
9. Center of PhenoGenomics, School of Life Sciences, EPFL; Swiss Federal Institute of Technology Lausanne, CH
10. Max Planck Institute of Molecular Cell Biology and Genetics, Dresden, Germany
11. Department of Molecular and Cell Biology, University of California, Merced, CA, USA

[*]Equal contributors. Author order is alphabetical.

### 1 Predicting protein signal from MS2 signal

An MS2 signal is an approximation of the instantaneous mRNA production rate. Given an MS2 signal, we predict the downstream protein signal by assuming the following flow of information, from MS2 ($x$) to total mRNA ($m$) to total protein ($p$)

$$x = \text{given} \tag{1}$$
$$\dot{m} = r_m x - \gamma_m m \tag{2}$$
$$\dot{p} = r_p m - \gamma_p p. \tag{3}$$

Here, the $r$'s are production rates, and $\gamma$'s are decay rates corresponding to mRNA and protein. Given an MS2 signal and this system of equations, the predicted protein is obtained by forward integration.

One of the challenges in integrating this system of equations is that the initial conditions for mRNA and protein are unknown. Here, we show that the effect of the initial condition on the predicted protein signal diminishes exponentially with a rate approximately equal to the protein decay rate. An example of this phenomenon is shown in Fig. A1 for a real MS2 trace. Therefore, to avoid artifacts from unknown initial conditions, we pick an arbitrary initial condition (usually zero mRNA and protein), but then throw out the predictions for mRNA and protein up to this timescale—we simply do not have the information to predict what those levels are.

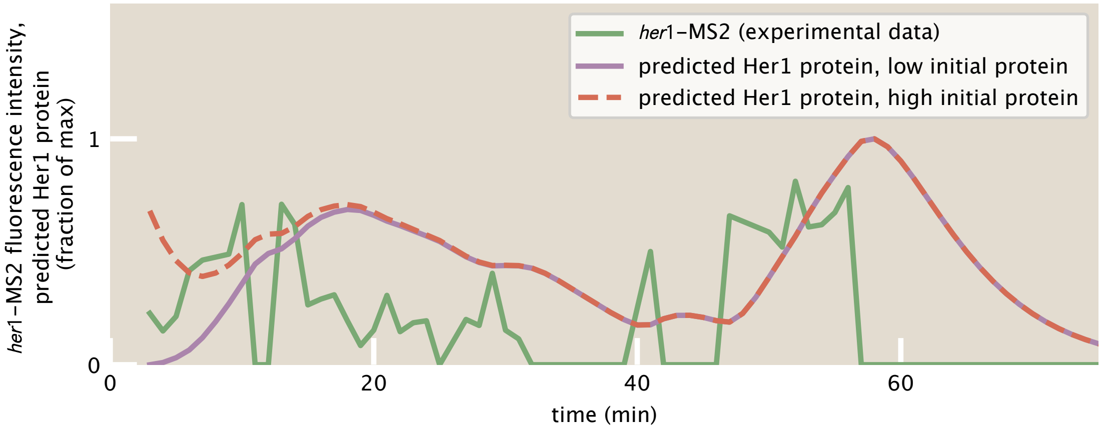


Figure A1: **Predicted protein traces for different initial conditions converge to the same solution**. In green we plot an experimental *her1*-MS2 fluorescence intensity trace. In purple is the predicted protein Her1 protein dynamics, assuming the the system starts with zero initial protein. For comparison, in red is the predicted Her1 protein dynamics starting from high initial protein. After approximately one oscillation cycle, the predicted protein dynamics become insensitive to the initial condition, which is unknown in our experiment. Parameters used are $\gamma_m = 0.23\ \text{min}^{-1}$, $\gamma_p = 0.23\ \text{min}^{-1}$. Each trace was rescaled to its maximum for visual clarity.

#### 1.1 Solving the system of equations using the Laplace transform

We show the effect of initial condition on the overall protein dynamics using the Laplace transform of the MS2 signal $x(t)$, which is given by

$$\mathcal{L}(x(t))(s) \equiv \tilde{x}(s) = \int_0^\infty dt\, \mathrm{e}^{-st} x(t). \tag{4}$$

Recall the relation between Laplace transform and differentiation:

$$\mathcal{L}\left(\frac{dx}{dt}(t)\right)(s) = s\tilde{x} - x(t=0). \tag{5}$$

Using this property, we can Laplace transform the equations for $m$ and $p$ above, leading to

$$s\tilde{m} - m(t=0) = r_m\tilde{x} - \gamma_m\tilde{m} \tag{6}$$
$$s\tilde{p} - p(t=0) = r_p\tilde{m} - \gamma_p\tilde{p}. \tag{7}$$

Let $m(t=0) \equiv m_0$ and $p(t=0) \equiv p_0$. Solving for $\tilde{m}(s)$ and $\tilde{p}(s)$ we find

$$\tilde{m}(s) = \frac{m_0}{s+\gamma_m} + r_m \frac{\tilde{x}}{s+\gamma_m} \tag{8}$$
$$\tilde{p}(s) = \frac{p_0}{s+\gamma_p} + r_p \frac{\tilde{m}}{s+\gamma_p} \tag{9}$$

The solution for each variable has 2 pieces, one coming from the initial condition, and one coming from the signal immediately upstream: $x$ for $m$, $m$ for $p$. We now need to invert the Laplace transform to solve for $m(t)$ and $p(t)$. A well-known Laplace transform-inverse transform pair is

$$\mathrm{e}^{-\gamma t} \Longleftrightarrow \frac{1}{s+\gamma}, \tag{10}$$

where the double arrow indicates the Laplace transform to the right and inverse Laplace transform to the left. With this result, the first term for $\tilde{m}(s)$ in equation 8 gives the exponential

$$\mathcal{L}^{-1}\left(\frac{m_0}{s+\gamma_m}\right) = m_0\mathrm{e}^{-\gamma_m t}. \tag{11}$$

The second term in equation 8 can be viewed as a product of two terms, $(s+\gamma_m)^{-1}$ and $\tilde{x}$. Like in Fourier space, in Laplace space the product of two Laplace transforms corresponds to convolution in real space. So, the inverse transform of this product is the convolution of the inverse transforms of each of the terms, resulting in

$$\mathcal{L}^{-1}\left(r_m \frac{\hat{x}}{s+\gamma_m}\right) = r_m\mathrm{e}^{-\gamma_m t} \star x(t). \tag{12}$$

Putting these together we have the solution

$$m(t) = m_0\mathrm{e}^{-\gamma_m t} + r_m\mathrm{e}^{-\gamma_m t} \star x(t). \tag{13}$$

To be specific, the convolution term is actually an incomplete convolution, in that it only goes up to time $t$, namely

$$m(t) = m_0 \mathrm{e}^{-\gamma_m t} + r_m \int_0^t dt' x(t') \mathrm{e}^{-\gamma_m (t-t')}. \tag{14}$$

This upper limit reflects the causality of the differential equation, i.e., mRNA at time $t$ is only determined by MS2 signal up to time $t$.

Analogously, the solution for $p(t)$ is

$$p(t) = p_0 \mathrm{e}^{-\gamma_p t} + r_p \int_0^t dt' m(t') \mathrm{e}^{-\gamma_p (t-t')}. \tag{15}$$

Plugging in the solution for $m(t)$, we get

$$p(t) = p_0 \mathrm{e}^{-\gamma_p t} + m_0 r_p \frac{1}{\gamma_m - \gamma_p} \left( \mathrm{e}^{-\gamma_p t} - \mathrm{e}^{-\gamma_m t} \right) + r_m r_p \int_0^t dt' \int_0^{t'} dt'' x(t'') \mathrm{e}^{-\gamma_m (t'-t'')} \mathrm{e}^{-\gamma_p (t-t')}. \tag{16}$$

The third term contains convolutions with decaying exponentials of rates given by the molecular decay rate $\gamma_m$ and $\gamma_p$. This part reflects the low-pass filtering done by the process of accumulating and destroying mRNA and protein. The first two terms reflect the initial condition, and we see explicitly that these terms decay to zero exponentially on the timescale set by the decay rates.

For the segmentation clock, the mRNA and protein decay rates are approximately equal. Setting them equal here, we simplify to

$$p(t) = p_0 \mathrm{e}^{-\gamma_p t} + m_0 r_p t \mathrm{e}^{-\gamma_p t} + r_m r_p \int_0^t dt' \int_0^{t'} dt'' x(t'') \mathrm{e}^{-\gamma_p (t-t'')}. \tag{17}$$

Thus, as $t \gg 1/\gamma_p$, the solution converges to a function that is independent of the initial condition, as shown in Fig. A1.

### 2 MS2 trace uncertainty

#### 2.1 Uncertainty due to background fluctuations

We quantify spot intensity by finding the spot center, estimating the background intensity levels around the spot, subtracting this background from each spot pixel (enforcing non-negativity), and then summing the pixels contained in an area defined by the spot size (i.e., by the point spread function).

Mathematically, the total spot intensity, $I$, is given by a sum of individual pixel intensities, $I_j$ minus the background, $B$:

$$I = \sum_{j \in \mathcal{V}} (I_j - B), \tag{18}$$

where the sum is over pixels contained in a neighborhood volume $\mathcal{V}$ around the spot center. Let there be $N$ pixels in $\mathcal{V}$. Then we can write $I$ as

$$I = \left( \sum_{j=1}^{N} I_j \right) - NB. \tag{19}$$

If we assume that the fluctuations in each pixel level, $I_j$, are all independent from one another (as is the typical model of shot noise in a microscope), and are independent from the background level, $B$, then the uncertainty in our measurement of the total spot intensity, $I$, is given by,

$$\sigma_I^2 = \sum_{j=1}^{N} \sigma_{I_j}^2 + N^2 \sigma_B^2. \tag{20}$$

The assumption that is typically made [1] is that the $\sigma_{I_j}$s are all the same and are equal to $\sigma_B$: that there is one typical scale of fluorescence fluctuations that affect the background and signal identically. Given this assumption, we then estimate $\sigma_B$ by looking at the fluctuations in the background. This is done by fitting a smooth trend to the time trace of the background signal (where the background level is obtained, for instance, by Gaussian fitting). A representative example of this background over time, together with its smooth polynomial fit, is shown in Fig. A2A. Then, the RMS deviation of the background from this smooth trend is measured, resulting in the error bars shown in Fig. A2B for non-zero *her1*-MS2 spot fluorescence intensities; the error bars on points with a value of zero come from spot detection uncertainties, as described below). Through 5-fold cross-validation, we found that fitting a polynomial of degree 4 to the background trend gives the lowest root-mean-squared-error fit without over-fitting (Fig. A2C).

With this estimate of $\sigma_B$, we get

$$\sigma_I^2 = \sum_{j=1}^{N} \sigma_B^2 + N^2 \sigma_B^2 = \sigma_B^2 (N + N^2). \tag{21}$$

Rearranging, we arrive at

$$\sigma_I = \sigma_B N \left( 1 + \frac{1}{N} \right)^{1/2}. \tag{22}$$

We use this equation to assign uncertainty to each spot fluorescence measurement.

#### 2.2 Including detection uncertainty

If we are relying on automated spot detection, there is also some additional uncertainty due to the probabilities of false positive and false negative detections. Here we incorporate

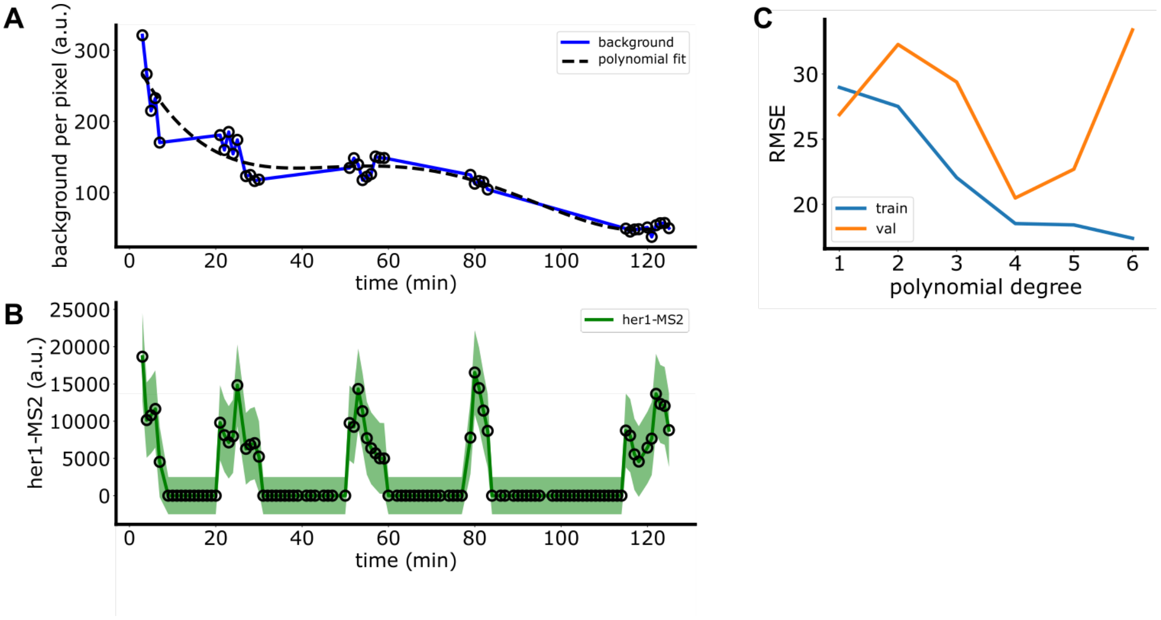


Figure A2: **Example of background fluctuation measurement.** (A) The mean background per pixel plotted over time for a given trace. Dashed line shows a polynomial fit of degree 4. (B) *her1*-MS2 trace with error bars derived from the background fluctuations as well as spot detection. (C) Root-mean-squared-error (RMSE) of the polynomial fit from (A) in a 5-fold cross-validation scheme, showing the train (blue) and validation (orange) errors. The minimum validation error occurs at polynomial degree 4.

those sources of uncertainty. The approach is to define a compound stochastic process that combines the probability of false positive/false negative detections with the measurement uncertainty obtained above.

Let the false positive probability by $f_p$ and the false negative probability be $f_n$. These are false positive and false negative rates for a whole dataset, measured by comparing a subset of traces to manually obtained traces, i.e. ground truth for the presence/absence of a spot. We break up the analysis into the case where we are adding uncertainty to detected spots and the case where we are adding uncertainty to a "zero" (no spot).

##### Case 1: a spot is detected

Suppose we detect a spot and measure its intensity $I$ and uncertainty $\sigma_I$. There is a probability $f_p$ that the spot is a false detection, and that the true intensity of the trace at this time point is zero. We want to incorporate this uncertainty.

Define a binary variable

$$\eta = \begin{cases} 1 & \text{if the spot is real (probability } 1 - f_p) \\ 0 & \text{if the spot is a false positive (probability } f_p). \end{cases} \tag{23}$$

Now define a stochastic variable $X$ that combines the intensity $I$, which we treat as a random variable, and the hidden "truth" variable $\eta$:

$$X = \begin{cases} I & \text{if } \eta = 1 \\ 0 & \text{if } \eta = 0. \end{cases} \tag{24}$$

The full uncertainty in our measurement of the intensity at this time point is the standard deviation of $X$. To calculate this uncertainty, we use the law of total variance, which states that

$$\text{Var}X = \mathbb{E}[\text{Var}(X|\eta)] + \text{Var}(\mathbb{E}[X|\eta]). \tag{25}$$

The first term, the average variance, is computed as follows. If $\eta = 1$, $X = I$, and $\text{Var}X = \sigma_I^2$. If $\eta = 0$, $X = 0$, and $\text{Var}X = 0$ (the intensity of a 'no spot' is exactly 0). The average variance is then given by

$$\begin{aligned} \mathbb{E}[\text{Var}(X|\eta)] &= \text{Var}(X|\eta = 1)P(\eta = 1) + \text{Var}(X|\eta = 0)P(\eta = 0) \\ &= \sigma_I^2(1 - f_p) + 0 * f_p = \sigma_I^2(1 - f_p), \end{aligned} \tag{26}$$

where in the first line $P(\eta = 1)$ is the probability that $\eta = 1$, which is $1 - f_p$, and analogously for the $\eta = 0$.

The second term, the variance of the averages, is similarly reasoned: if $\eta = 1$, $X = I$, and given that we only have one measurement of $I$, we assume $\mathbb{E}[X] = I$. If $\eta = 0$, $X = 0$, and $\mathbb{E}[X] = 0$. For convenience, define $M := \mathbb{E}[X|\eta]$. The average of $M$ is given by

$$\bar{M} = I(1 - f_p) + 0 * f_p = I(1 - f_p). \tag{27}$$

Now we can compute

$$\begin{aligned} \text{Var}(\mathbb{E}[X|\eta]) &= \mathbb{E}[(M - \bar{M})^2] \\ &= (I - I(1 - f_p))^2(1 - f_p) + (0 - I(1 - f_p))^2 p \\ &= I^2 f_p^2(1 - f_p) + I^2(1 - f_p)^2 f_p \\ &= I^2 f_p(1 - f_p)(f_p + 1 - f_p) \\ &= I^2 f_p(1 - f_p). \end{aligned} \tag{28}$$

Putting these two results together, we arrive at

$$\text{Var}X = \sigma_I^2(1 - f_p) + I^2 f_p(1 - f_p), \tag{29}$$

or

$$\sigma_X = \sqrt{\sigma_I^2(1 - f_p) + I^2 f_p(1 - f_p)}. \tag{30}$$

##### Case 2: no spot is detected

Here, we measure a zero, but there is a probability $f_n$ that we missed a true spot with non-zero intensity. We will estimate the "missing" intensity by the empirical mean of the trace, $\bar{I}$—an exact number, not a stochastic one.

The approach here mirrors case 1, but is simpler, because the "missing" value is exact. Define

$$\eta = \begin{cases} 1 & \text{if there is really no spot (probability } 1 - f_n) \\ 0 & \text{if we missed a spot (i.e. false negative with probability } f_n). \end{cases} \tag{31}$$

In this case, $\mathrm{Var}(X|\eta) = 0$ for both values of $\eta$, so we simply have

$$\mathrm{Var}X = \mathrm{Var}(\mathbb{E}[X|\eta]). \tag{32}$$

Let $M := \mathbb{E}[X|\eta]$. Its average

$$\bar{M} = \bar{I} f_n. \tag{33}$$

Then we have

$$\begin{aligned} \mathrm{Var} &= (\bar{I} - \bar{I} f_n)^2 f_n + (\bar{I} f_n)^2 (1 - f_n) \\ &= \bar{I}^2 (1 - f_n)^2 f_n + \bar{I}^2 f_n^2 (1 - f_n) \\ &= \bar{I}^2 f_n (1 - f_n)(1 - f_n + f_n) \\ &= \bar{I}^2 f_n (1 - f_n). \end{aligned} \tag{34}$$

In terms of the final uncertainty,

$$\sigma_X = \bar{I} \sqrt{f_n (1 - f_n)}. \tag{35}$$

##### Conclusion

Putting all these results together, we arrive at the final uncertainty of our measurement

$$\sigma = \begin{cases} \sqrt{\sigma_I^2 (1 - f_p) + I^2 f_p (1 - f_p)} & I > 0 \\ \\ \bar{I} \sqrt{f_n (1 - f_n)} & I = 0 \end{cases} \tag{36}$$

where $\sigma_I$ is the uncertainty from background fluctuations, $f_p$ is the false-positive rate of detection, $f_n$ is the false-negative rate of detection, $I$ is the intensity of the individual spot, and $\bar{I}$ is the mean intensity of the trace.

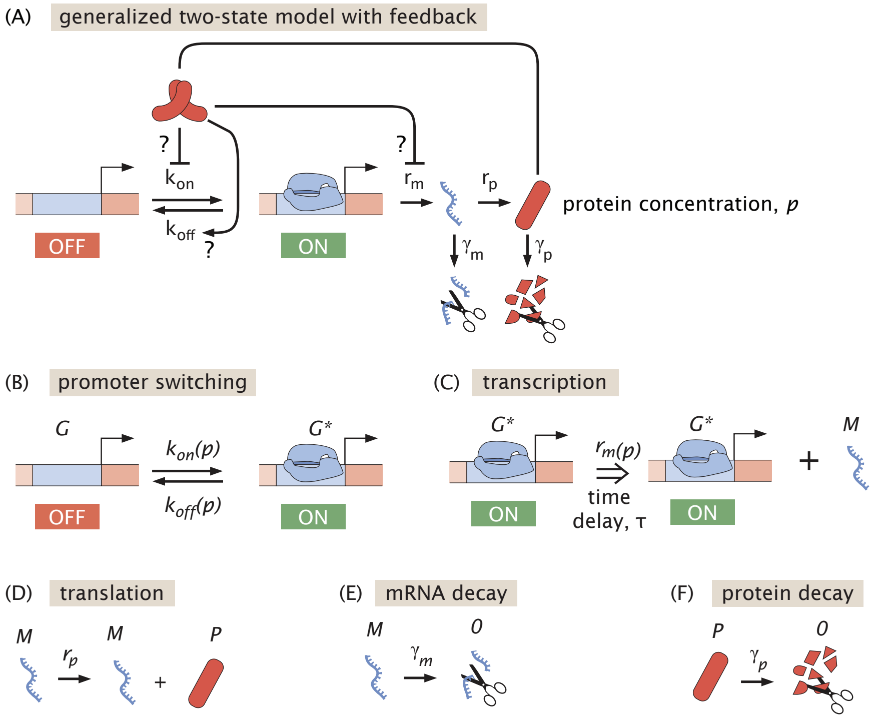


Figure A3: **Overview of the bursting model.** (A) Cartoon of the overall model structure. The *her1* gene switches between ON and OFF states. In the ON state, mRNA is produced, which is translated into protein. The Her1 protein has the ability to regulate the ON-OFF switching rates and/or the transcription rate. Both mRNA and proteins also decay. (B)-(F) Cartoons showing the individual reactions that make up the model: (B) gene state switching; (C) transcription, which is a delayed reaction that takes a time $\tau$ to complete; (D) translation; (E), mRNA decay; (F) protein decay. In these cartoons, variables in capital letters—$G$, $G^*$, $M$, and $P$—represent individual molecules, while lowercase $p$ denotes the concentration of proteins.

### 3 Generalized transcriptional bursting model with feedback

In this section we provide details on the mathematical model from Fig. 6 of the main text. The goal of the model is to provide a minimal phenomenological description of transcriptional bursting with auto-repression that is flexible enough to include a range of possible regulatory mechanisms.

##### 3.1 Statement of the model

The model is fully stochastic and is described by the following set of chemical reactions:

$$G \underset{k_{off}(p)}{\overset{k_{on}(p)}{\rightleftarrows}} G^* \tag{37}$$

$$G^* \overset{r_m(p)}{\Longrightarrow_\tau} G^* + M \tag{38}$$

$$M \xrightarrow{r_p} M + P \tag{39}$$

$$M \xrightarrow{\gamma_m} \emptyset \tag{40}$$

$$P \xrightarrow{\gamma_p} \emptyset \tag{41}$$

The variables $G$ and $G^*$ represent the OFF and ON promoter states, respectively. The variables $M$ and $P$ schematically represent individual mRNA and protein molecules, respectively, while $p$ denotes the number of $P$ molecules. The rates $k_{on}$, $k_{off}$, and $r_m$ are generically functions of $p$ and encode the types of autoregulation that occur. The double arrow notation $\Longrightarrow_\tau$ denotes a delayed reaction that takes a time $\tau$ to complete. Computationally, the delayed transcription reaction is initiated with a rate the depends on the current protein concentration, but the new mRNA is not added to the system until a time $\tau$ later, following [2]. This time delay is crucial for generating oscillations under the amplitude regulation scheme, but is not necessary for frequency or duration regulation schemes. Thus, the general model is a delayed stochastic system that is simulated using the delayed Gillespie algorithm of Bratsun et al. [2] implemented in Python.

**Amplitude regulation:**

We model amplitude regulation via a Hill function

$$r_m(p) = r_{m,0} + r_{m,1}\frac{1}{1 + (p/K_D)^n}. \tag{42}$$

The transcription rate has a basal rate, $r_{m,0}$, and a protein-dependent rate with prefactor $r_{m,1}$. The protein-dependence is described by a decreasing Hill function with coefficient, $n$, and scale $K_D$. For simplicity, we set $r_{m,0} = 0$. Obtaining robust oscillations under the amplitude regulation model requires addition time delay to be introduced into the system. We do this by explicitly adding a time delay, $\tau$, between the initiation of the mRNA production reaction and its completion within the simulation [2]. In this model, the rate of transcription initiation depends on the current number of Her1 proteins, but the time required to produce mature *her1* mRNAs is modeled explicitly.

Under amplitude regulation, switching between $G$ and $G^*$ is not required to generate oscillations. Rather, the sharp modulation of the transcription rate by Her1 proteins on their own can produce burst-like behavior.

**Frequency regulation:**

We model frequency regulation via

$$k_{on} = k_{+,0} + k_{+,1} \frac{1}{1 + (p/K_D)^n}. \tag{43}$$

In this model, Her1 protein decreases the rate at which the promoter switches from OFF to ON. For simplicity, we set $k_{+,0} = 0$. Also for simplicity, we ignore the time delay in mRNA production, since it is not required to generate quasi-periodic solutions (i.e., we set the time delay $\tau = 0$).

**Duration regulation:**

We model duration regulation via

$$k_{off} = k_{-,0} + k_{-,1} \frac{p^n}{K_D^n + p^n}. \tag{44}$$

In this model, Her1 protein increases the rate at which the promoter switches from ON to OFF. For simplicity, we set $k_{-,0} = 0$. Also for simplicity, we ignore the time delay in mRNA production, since it is not required to generate quasi-periodic solutions (i.e., we set the time delay $\tau = 0$).

#### 3.2 Connection to deterministic models of the segmentation clock

The standard approach to modeling segmentation clock oscillations [3], which predicts smooth mRNA oscillations, is to use a set of delay differential equations describing mRNA and protein concentrations, $m$ and $p$, that is of the form

$$\dot{m} = r_m f(p(t-\tau)) - \gamma_m m \tag{45}$$
$$\dot{p} = r_p m - \gamma_p p. \tag{46}$$

Here, $r_m$ and $r_p$ are the transcription and translation rates of mRNA and protein, respectively, and $\gamma_m$ and $\gamma_p$ are the decay rates of mRNA and protein, respectively. The function $f(p)$ describes auto-regulation, with the rate of transcription modulated by protein concentration. This function typically takes the form of a decreasing Hill function,

$$f(p) = \frac{1}{1 + (p/K_D)^n}. \tag{47}$$

The time delay in the repression term is required mathematically for the system to contain a Hopf bifurcation [3, 4]. Past the Hopf bifurcation, this dynamical system exhibits oscillatory solutions, with a period set by the sum of the delay and the molecular half-lives [3, 4].

We can obtain this deterministic model as a particular limit of our stochastic bursting model. The first step to obtain the deterministic limit is to consider the amplitude regulation scheme in the limit where transcription and translation rates are high, such that small number fluctuations in mRNA and protein concentrations are negligible and we can approximate them by continuous variables that evolve deterministically. In that limit, the system becomes a piece-wise deterministic process,

$$\dot{m} = r_m \chi_t \frac{1}{1 + \left(\frac{p(t-\tau)}{KD}\right)^n} - \gamma_m m \tag{48}$$

$$\dot{p} = r_p m - \gamma_p p, \tag{49}$$

with $\chi_t$ the promoter state variable that stochastically switches between ON and OFF states with constant probability rates $k_{on}$ and $k_{off}$. Then, we simply take the limit of an always on promoter, $k_{on}/k_{off} \to \infty$, such that the probability that $\chi_t = 1$ limits to unity. In this limit, we recover the deterministic model.

#### 3.3 Connection to other work

Our model contains as special cases models from various other works. In particular, Wang et al.[5] studied a protein-only version of the duration regulation model in the case of linear repression, $k_{off} = k_{-,1}p$, and found approximate analytic conditions for quasi-regular oscillations. However, the protein oscillations in this linear model are not as regular as experimentally-measured protein oscillations; we found that non-linearity appears to help achieve cleaner oscillations.

Work by Zinani et al. [6] considers a fully stochastic amplitude regulation model, using a high burst frequency of approximately 4 / minute. As we imaged every 1 minute in our experiments, we cannot rule in or out dynamics at that fast of a timescale.

The model of Lengyel et al. [7] does not assume bursting in the absence of auto-repression as our model allows, but posits a mechanism similar to our version of amplitude regulation. By explicitly modeling the binding of Her1 proteins to the *her1* regulatory DNA, they demonstrated that regular oscillations can arise without incorporating additional delays into the system.

The model of Jenkins et al. [8] considers explicit binding and unbinding of Her1/7 dimers to a single binding site and two copies of each gene within an more complex gene regulatory network. When the model is restricted to only one Her species, it resembles our version of duration regulation, with unbounded regulation of the promoter off rate, $k_{off} \sim p^2$.

Finally, there is a large literature on stochastic oscillations in the circadian clock, including Bratsun et al. [2], which is a more restricted version of our model.
